# Negative allosteric modulation of the glucagon receptor by RAMP2

**DOI:** 10.1101/2022.08.30.505955

**Authors:** Kaavya Krishna Kumar, Evan S. O’Brien, Chris H. Habrian, Naomi R. Latorraca, Haoqing Wang, Inga Tuneew, Elizabeth Montabana, Susan Marqusee, Daniel Hilger, Ehud Y. Isacoff, Jesper Mosolff Mathiesen, Brian K. Kobilka

**Affiliations:** Department of Molecular and Cellular Physiology, Stanford University School of Medicine, 279 Campus Drive, Stanford, CA 94305, USA; Department of Molecular and Cell Biology, University of California, Berkeley, CA 94720; Zealand Pharma A/S, Sydmarken 11, Soborg 2860, Denmark; QB3 Institute for Quantitative Biosciences, University of California, Berkeley, CA 94720; Department of Chemistry, University of California, Berkeley, CA 94720; Department of Pharmaceutical Chemistry, Philipps-University Marburg, Marcher Weg 6, Marburg 35037, Germany; Helen Wills Neuroscience Institute, University of California, Berkeley, CA, 94720, USA

## Abstract

Receptor activity-modifying proteins (RAMPs) modulate the activity of many Family B heterotrimeric guanine nucleotide-binding protein (G protein)-coupled receptors (GPCRs). The glucagon receptor (GCGR), a Family B GPCR responsible for maintenance of proper blood sugar levels, interacts with RAMP2, though the purpose and consequence of this interaction is poorly understood. Using a series of biochemical and cell-based assays, we show that RAMP2 interacts with and broadly inhibits GCGR-induced downstream signaling. Hydrogen-deuterium exchange monitored by mass spectrometry (HDX-MS) demonstrates that RAMP2 enhances local flexibility in select locations in and near the receptor extracellular domain (ECD) as well as at a key region in the 6^th^ transmembrane helix, while single-molecule fluorescence resonance energy transfer (smFRET) experiments show that this RAMP2-induced ECD disorder results in inhibition of active and intermediate states of the intracellular face of the receptor. Using cryo-electron microscopy (cryoEM), we determined the structure of the GCGR-G_s_ complex at 2.9 Å resolution in the presence of RAMP2. RAMP2 apparently does not interact with GCGR in an ordered manner, yet the ECD of GCGR is indeed largely disordered in the presence of RAMP2. This disorder is accompanied by rearrangements of several key areas of the receptor, resulting in the formation of a likely unproductive complex. Together, our studies suggest that RAMP2 acts as a negative allosteric modulator of GCGR by enhancing conformational sampling of the ECD.

## Introduction

The glucagon receptor (GCGR) is a Family B heterotrimeric guanine nucleotide-binding protein (G protein)-coupled receptor (GPCR) responsible for maintaining proper blood glucose concentrations^1^. Glucagon binding to GCGR activates the receptor, resulting in cyclic adenosine monophosphate (cAMP) production predominantly via GCGR interaction with the adenylyl cyclase stimulatory G protein, G_s_^2^. We previously demonstrated that Family B GPCRs, like GCGR, are not as effective at activating G_s_ as their Family A counterparts, likely due to the kinetically-limiting necessity of a large helical break that occurs in TM6 upon Family B GPCR activation^3^. We were therefore interested in characterizing modulatory factors that could stimulate this activity. Receptor activity-modifying proteins (RAMPs) are single-pass TM proteins with an N-terminal extracellular domain that bind Family B GPCRs and modify their ligand binding and intracellular signaling^4,5^. These effects are exemplified by the structural and dynamic changes induced by binding of one of the three RAMPs to either the calcitonin receptor (to form the amylin receptors)^6^ or the calcitonin receptor-like receptor (to form the calcitonin gene-related peptide receptor or adrenomedullin receptors)^7,8^. However, the effects of RAMP interaction with other classes of Family B receptors are less well understood. RAMP2 is known to interact with GCGR^9^ resulting in a disparate array of ligand binding and intracellular consequences, including increased glucagon potency^10^, decreased potency^11^ decreased cAMP production over a prolonged period^12^ or cAMP accumulation^10^. To address these disparate findings, we purified monomeric RAMP2 in the absence of receptor and performed a variety of biochemical, biophysical and structural studies to more directly characterize the effect of non-constitutive RAMP2 binding on GCGR activity.

## Results

Our previous study showed that GCGR (along with other Class B GPCRs) is significantly slower at facilitating nucleotide exchange in G_s_ compared to Class A GPCRs^3^. We initially speculated that RAMP2 may serve as a cofactor (positive allosteric modulator) to increase GCGR induced G_s_ turnover. To understand how RAMP2 binding impacts GCGR signaling, we purified monomeric RAMP2 and conducted a time-dependent in vitro GTPase-turnover assay (Fig. S1A)^13^. While agonist-bound GCGR demonstrates guanine nucleotide exchange factor (GEF) activity that is very weakly sensitive to the initial concentration of GDP ([GDP]_i_) (Fig. 1A, Fig. S1B), pre-incubation with RAMP2 (2:1 RAMP2:GCGR) results in potent inhibition of GEF activity in a [GDP]_i_ dependent manner. This inhibitory effect is present at all time points over the course of the nucleotide depletion assay (Fig. S1B). RAMP2 has no effect on GCGR-mediated G_s_ turnover in the absence of GDP, and no impact on basal G_s_ turnover (data not shown) or turnover stimulated by agonist-bound β_2_-adrenergic receptor (β_2_AR, a Family A GPCR) (Fig. S1C), suggesting that the mechanism of inhibition is through specific interaction with GCGR. Because of the [GDP]_i_-dependent nature of RAMP2 inhibition, we hypothesized that RAMP2 acts by inhibiting receptor-catalyzed nucleotide release from G_s_. Consistent with this mechanism of inhibition, pre-incubation of GCGR with RAMP2 decreases the observed Gαs ^3^H-GDP dissociation rate (*k*_off_) from 0.0033 s^−1^ to ~0.0001 s^−1^ (Fig. 1B). Thus, contrary to our initial speculation, RAMP2 appears to potently *inhibit* GCGR signaling through G_s_ by inhibiting the initial GDP release step; this manifests in the GTP turnover assay as a GDP-titratable inhibitory effect. Because the GTP-depletion rate of GCGR in the presence of RAMP2 is even *slower* than intrinsic G_s_ activity alone (Fig. 1A, S1B), we speculated that GCGR/RAMP2 heterodimers may act as a sink for GDP-bound G_s_. Indeed, while addition of agonist-bound GCGR to β_2_AR has little impact on turnover by the β_2_AR, co-incubation with RAMP2/GCGR/agonist with β_2_AR results in significant inhibition of β_2_AR-mediated turnover, suggesting that GCGR/RAMP2 acts to actively sequester G_s_ away from other G_s_-coupled receptors (Fig. S1D).

**Fig 1.**
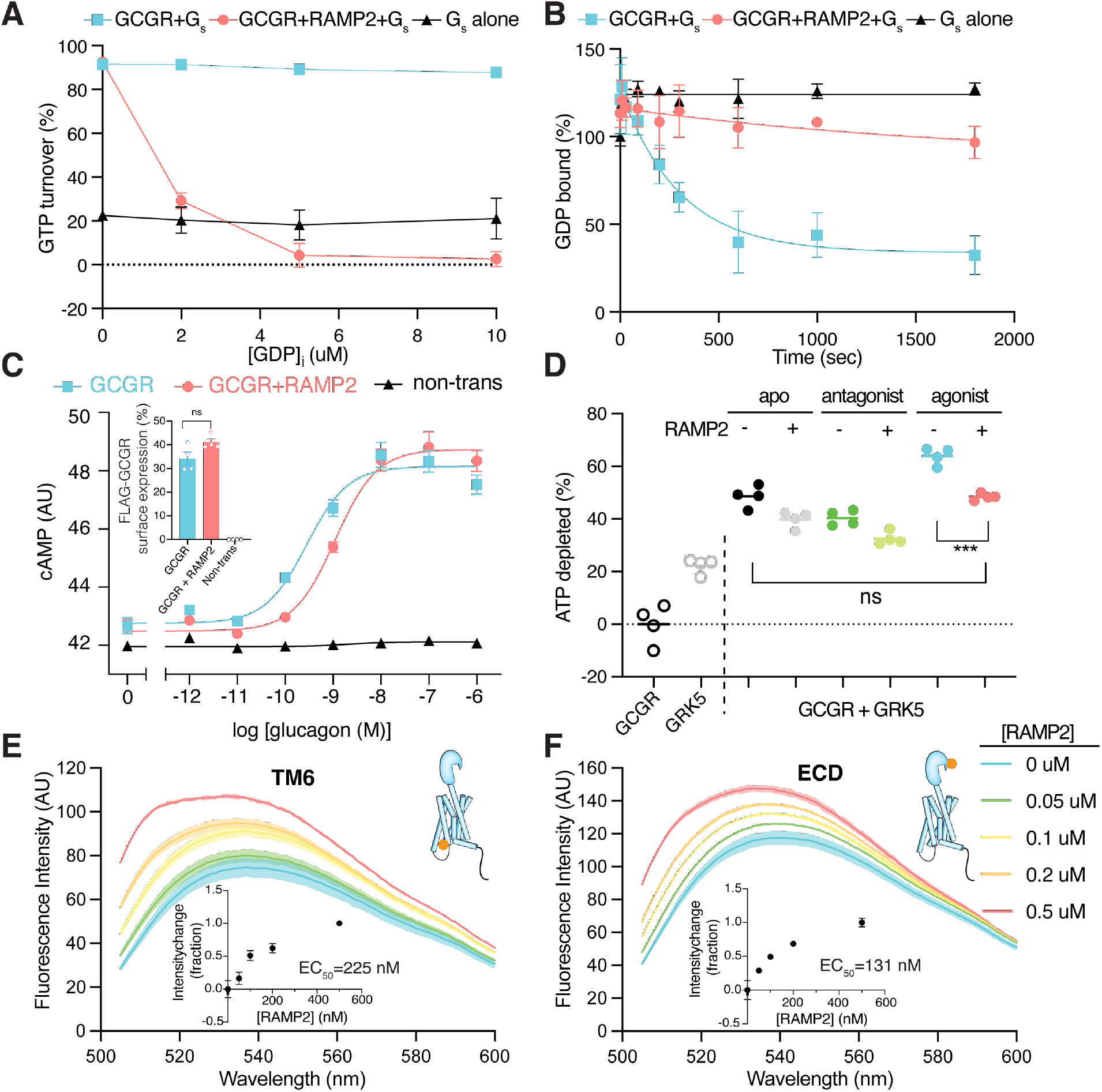
Functional consequences of the widespread perturbations induced in GCGR upon interaction with RAMP2. (**A**) The GTP turnover assay shows that GCGR activation of G_s_ is largely independent of GDP concentration, but upon pre-coupling of GCGR with RAMP2, G_s_ activation is potently inhibited in a [GDP] dependent manner and appears to inhibit even intrinsic G_s_ activity. (**B**) The rate of ^3^H-GDP release from G_s_ is significantly reduced when GCGR is pre-incubated with RAMP2. (**C**) Cells transfected with GCGR display increases in cAMP levels upon stimulation with glucagon with an observed EC_50_ of 0.28 nM (pEC_50_=9.6 [95% confidence limits 9.8-9.3]); cells expressing the same levels of GCGR in the presence of excess transfected RAMP2 display an EC_50_ of 1.10 nM (pEC_50_=9.0 [95% confidence limits 9.1-8.8]), a 4-fold right-shift in potency. (**D**) Using an ATP depletion assay, we show that the presence of RAMP2 inhibits receptor phosphorylation regardless of what is bound in the orthosteric site, though the inhibitory effect is most significant for agonist-induced phosphorylation, where receptor phosphorylation returns to levels seen in the apo-state. Using agonist-bound GCGR site-specifically labeled at the intracellular end of TM6 (**E**) or on the N-terminal helix of the ECD (**F**) with the environmentally sensitive fluorophore NBD, we show that addition of increasing amounts of RAMP2 results in titratable environmental changes in both regions of the receptor.

Previous studies interrogating the effect of RAMP2 on GCGR-G_s_ signaling with mixed results have been performed with different cell types and/or potentially different RAMP2-receptor ratios^10–12^. Since overexpression of GCGR (but not β_2_AR) leads to a concurrent increase in RAMP2 surface expression (Fig. S1E), we carefully optimized the surface-expression levels of GCGR and RAMP2 for cellular signaling assays in HEK293 cells. After selecting transfection conditions for similar surface expression levels of GCGR without and with co-expression of RAMP2, we performed a non-IBMX perturbed cyclic adenosine monophosphate (cAMP) assay as a direct measure of GCGR-G_s_ signaling. At similar surface expression levels of GCGR expression, the presence of RAMP2 results in a significant decrease in glucagon potency EC_50_= 0.28 nM vs 1.10 nM) (Fig. 1C), consistent with the G_s_ inhibition observed with biochemical experiments.

In addition to G proteins, ligand-activated Family B GPCRs have been shown to signal through β-arrestins upon being phosphorylated by GPCR kinases (GRKs)^12,14–16^. Multiple studies have shown that RAMP2 decreases β-arrestin recruitment to GCGR^11,12^. To understand if this decrease in β-arrestin coupling is due to the role of RAMP2 in modulating GRK phosphorylation of GCGR, we measured GRK-mediated GCGR phosphorylation in the presence and absence of RAMP2 with ligands of different efficacy, including a newly engineered, more soluble antagonist peptide devoid of apparent intrinsic efficacy (Fig. S2), using a luciferase-based phosphorylation assay (Fig. S3A). Compared to unliganded (apo)-receptor, phosphorylation by GRK5 is reduced in antagonist-bound GCGR and increased by our previously developed soluble agonist peptide (Fig. 1D)^3^. RAMP2 broadly diminishes GRK5-phosphorylation irrespective of the ligand state of the receptor, aside from the negative allosteric modulator L-168,049 (Fig. 1D; Fig. S3B). This effect is largest for the agonist-bound form of GCGR, where RAMP2 results in GRK5 phosphorylation levels similar to the apo-form of the receptor (Fig. 1D). In-gel staining for phosphorylated protein further confirmed that RAMP2 dramatically inhibits GRK5-mediated phosphorylation of GCGR (Fig. S3C). However, GRK2 phosphorylation of GCGR is not impaired by the presence of RAMP2 (Fig. S3D) suggesting that RAMP2 specifically inhibits phosphorylation of GCGR by certain GRK isoforms.

To probe the impact of RAMP2 binding on the conformation(s) of GCGR, we employed fluorescence spectroscopy, wherein we site-specifically labeled the receptor on an introduced cysteine at the cytoplasmic end of TM6 (F349C) and the ECD (K31C) with the environmentally sensitive fluorophore 4-chloro-7-nitrobenz-2-oxa-1,3-diazole (NBD) in a minimal cysteine GCGR (mC GCGR) background^3^. Incremental addition of RAMP2 to agonist-bound mC-GCGR-349C-NBD results in a titratable (EC_50_~225 nM) increase in fluorescence (Fig. 1E), consistent with increased occupancy of the inactive conformation of TM6^3^. This may provide an explanation for the decrease in activity observed in the GTP turnover assay. In addition to perturbing the intracellular conformational distribution of GCGR, RAMP2 *also* induces significant changes in fluorescence of the ECD sensor (Fig. 1F, EC_50_~131 nM). Taken together, these experiments demonstrate that RAMP2 interaction with GCGR induces widespread conformational changes in the receptor from the ECD to the intracellular face of the receptor.

Next, we performed hydrogen-deuterium exchange monitored by mass spectrometry (HDX-MS) to quantify changes in local conformational flexibility across the receptor upon binding of RAMP2 (Fig. 2A, Fig. S4A,B). For HDX-MS we used antagonist-bound GCGR as our pull-down experiment showed RAMP2 interacts most stably with antagonist-bound GCGR (Fig. S4C). HDX-MS was performed on GCGR in the absence or presence of excess RAMP2. We obtained high-quality peptide coverage for 64.8% of the GCGR sequence, with most peptides reporting on the ECD, TM1, TM2, TM6, TM7, and the C-terminus (Fig. S4A,B). We also performed HDX-MS on RAMP2 alone. We obtained high-quality coverage of 41.1% of the RAMP2 sequence, with peptides primarily derived from the single-pass transmembrane segment (Fig. S4A,B). Peptides within the RAMP2 transmembrane segment, spanning residues 132–163, exhibited reduced deuterium uptake in the presence of GCGR (Fig. 2A,B). While not a quantitative measure of complex formation (due to the excess RAMP2 in the complex study), reduced exchange in this transmembrane segment is consistent with an *AlphaFold2*-predicted model^17^ of the GCGR–RAMP2 complex, in which the RAMP2 transmembrane segment forms an extended binding interface with TM3, TM4 and TM5 of GCGR (Fig. 2A).

**Fig 2.**
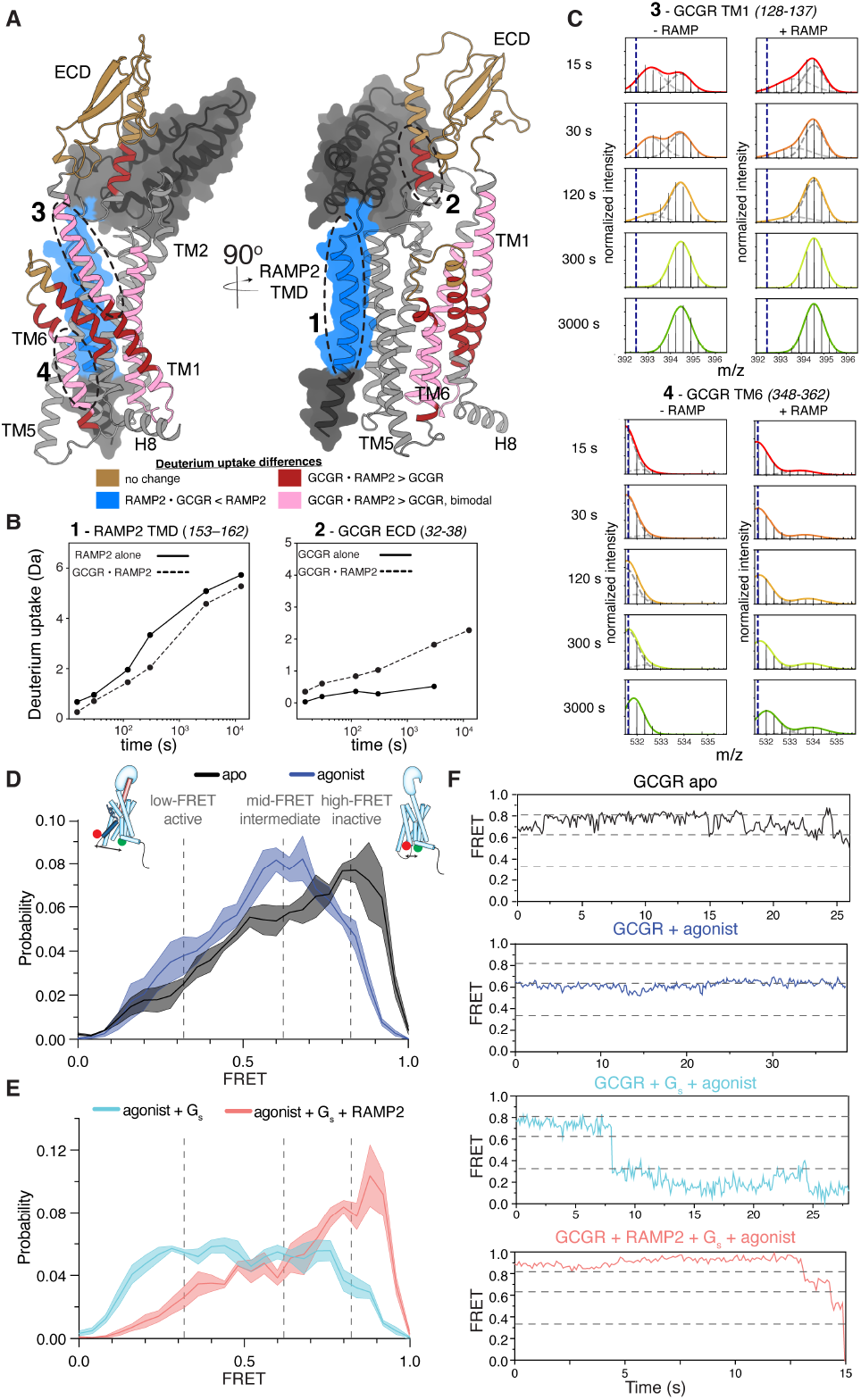
RAMP2 effects on GCGR local and global dynamic behavior. **(A)** *Alphafold* model of GCGR-RAMP2 complex onto which changes in HDX are plotted upon complex formation. **(B)** Example HDX-MS exchange curves in the presence and absence of excess RAMP2 show that several key regions of the RAMP2 (153-162) and receptor (32-28) are impacted by heterodimer complex formation. (**C**) HDX-MS plots showing bimodal distribution in GCGR TM1 (residues 128-137), an indication of conformational heterogeneity. This heterogeneity is increased in the presence of RAMP2. Though not present in the absence of RAMP2, a bimodal distribution is induced in TM6 (348-362) upon addition of RAMP2. (**D**) smFRET experiments show that a dominant high-FRET state (~0.83) of GCGR labeled with donor and acceptor fluorophores at the intracellular ends of TM4 and TM6 is present in the absence of orthosteric agonist (**D**, black) with brief excursions to mid-(~0.63) FRET (~0.32) states (**F**, black), and the binding of agonist peptide results in a dominant mid-FRET state at the expense of the high-FRET state (**D**,**F**; blue). G_s_ coupling to agonist-bound GCGR increases the population of the low-FRET, suggesting a full outward movement of TM6 at the expense of the mid-FRET agonist-specific intermediate state and high-FRET inactive state (**E**,**F**; cyan). However, pre-coupling of agonist-bound GCGR with RAMP2 potently inhibits a G_s_-induced increase in the population of the fully outward TM6 conformation as well as the agonist-associated intermediate state (**E**,**F**; salmon).

For GCGR in the presence of RAMP2, we found a surprising enhancement of deuterium uptake in the N-terminal region of the ECD, including a peptide corresponding to residues 32–38 (Fig. 2B). This region (32–38) is adjacent to the NBD-labeling position in the ECD, and the changes responsible for the increase in HDX in this region likely contribute to the NBD fluorescence change (Fig. 1F). Structural elements within the remainder of the ECD exhibited no difference in deuterium uptake upon RAMP2 binding (Fig. S4A), indicating that RAMP2 likely perturbs the ECD through a rigid-body motion and not by altering its local structure. Multiple studies have shown the importance of ECD dynamics in Family B receptor ligand binding kinetics, activation and signaling^18–21^.

In addition to the ECD, RAMP2 binding also resulted in complex hydrogen exchange behavior elsewhere in the receptor, indicating RAMP2-enhanced conformational heterogeneity. This effect is most easily seen in peptides that display bimodal behavior, both in terms of an increased number of peptides with bimodal behavior and by a change in the ratio of the two peaks for those that show bimodal behavior in the isolated receptor. Bimodal mass-isotope distributions are seen in peptides that include the N-terminal portion of TM1 and portions of TM2 and TM6 (Fig. 2C, Fig. S4A,B). Such bimodal behavior can arise when the rate of labeling is faster than the rate of interconversion between a closed, exchange-incompetent conformation and an open, exchange-competent conformation (EX1 or EXX kinetics)^22^. For example, RAMP2 binding increases population of the heavier (right-hand) peak for peptides covering the beginning of TM1 (residues 128–144) (Fig. 2C), suggesting that RAMP2 increases the rate of conversion between two conformations that TM1 populates within the native conformational ensemble of GCGR in the absence of RAMP2. While in CLR-RAMP structures (which are obligate dimers) TM1 is not in direct contact with RAMP2, in GCGR the extracellular end of TM1 might unravel to accommodate ECD conformational changes upon RAMP binding. Mutations in this region have been shown to decrease agonist peptide binding^23^, suggesting a possible functional link to RAMP2-induced changes in conformational flexibility. Additionally, within TM6, peptides containing the PXXG motif, display bimodal behavior only the presence of RAMP2 (Fig. 2C). The PXXG motif of TM6 has been shown to be pivotal for receptor activation^24^. Increased hydrogen exchange upon RAMP2 binding is not universal: other locations in GCGR, including in the C-terminus, do not exhibit changes in deuterium uptake upon RAMP2 binding (Fig. S4A), demonstrating that the increases in observed dynamics are not a result of broad-scale receptor destabilization. Taken together, our HDX-MS results show that RAMP2 selectively modulates the intrinsic conformational heterogeneity of several key structural regions within GCGR, including the ECD and TM6.

We next used single-molecule fluorescence resonance energy transfer (smFRET) to study the effect of RAMP2 on the conformational dynamics of the intracellular face of GCGR. Using our mC-GCGR construct, we site-specifically labeled a previously characterized pair of introduced cysteines, one in TM4 as a reference site (265C) and one in the activation-sensitive TM6 (345C)^3^ with cysteine-reactive versions of LD555 (donor) and LD655 (acceptor) fluorophores to probe TM6 outward movement upon ligand and RAMP2 binding (using nitroxide spin labels observed by cwEPR to characterize and minimize background labeling; see Fig. S5A for more details). The fluorescently-labeled mC-GCGR-265C/345C protein was immobilized at low density on a PEG-passivated coverslip with a biotinylated anti-FLAG antibody, and imaged using total internal reflection (TIRF) microscopy^25^. In the apo-state, GCGR exhibits a broad distribution of FRET efficiencies with a major peak centered at ~0.83 (Fig. 2D, black). This high-FRET state likely corresponds to the inactive, inward conformation of TM6, which brings the intracellular end of TM6 into close proximity to the intracellular end of TM4, as seen previously^3,13^. Representative individual traces of apo-GCGR largely occupy the high-FRET state with occasional transitions to a mid-FRET state with a FRET efficiency ~0.63 (Fig. 2F). Antagonist peptide or RAMP2 addition have a minimal effect on the FRET distribution relative to apo receptor (Fig. S5B), consistent with all three conditions representing a similar ensemble of inactive states. In the presence of full agonist peptide, heterogeneity in the distribution of FRET values remains, but the mid-FRET peak ~0.63 becomes dominant at the expense of the inactive, high-FRET state along with an increase in a low-FRET peak at FRET efficiency ~0.32 (Fig. 2D, dark blue). This mid-FRET peak appears to be an agonist-specific intermediate state of the receptor rather than an average of inactive/active states because we are able to resolve transitions to and from this state when imaging with a 100 ms frame rate^26^ (Fig. 2F; Fig. S5C). This is in apparent contradiction with previous DEER results suggesting that GCGR has no significant change in distance between TM4 and TM6 upon binding to glucagon, though some subtle but significant changes in the bimane fluorescence spectra upon agonist binding were observed^3^. We speculate that subtle changes in conformation at the intracellular end of TM6 that are not observable by the pure distance changes measured by DEER are amplified by the large, flexible dyes used for smFRET experiments, as well as the inherently complex nature of observed FRET efficiencies^27^.

Formation of a nucleotide-free GCGR-G_s_ complex results in a substantial increase in occupancy of the low-FRET state (~0.32) (Fig. 2E, 2F, cyan), suggesting that this state is the fully outward, active state of TM6 observed in the structure of GCGR/G_s_ complex. The FRET efficiency distribution does not shift completely to the active conformation of TM6. This may reflect either incomplete complexing efficiency of the receptor with G_s_, or intrinsic receptor dynamics within the G_s_-bound ensemble. Addition of RAMP2 to agonist-bound receptor results in a near complete elimination of fully-active and agonist-associated intermediate states of TM6 in favor of an inactive-like conformation (Fig. S5B). Further, the addition of RAMP2 to agonist-bound GCGR in the presence of G_s_ shifts occupancy back to the high-FRET inactive state that is comparable to, or even above, that of the apo-state (Fig. 2E, 2F, salmon) and interestingly, not to the agonist-bound intermediate state. Hence, by inhibiting the formation of the agonist-associated intermediate and fully active conformations of GCGR, RAMP2 binding decreases the probability of productive agonist-induced GCGR-G_s_ interaction, conceivably explaining at least part of the decreased GTP turnover (Fig. 1A) and GDP release rates (Fig. 1B).

The observation that GCGR/RAMP2 co-complexes can suppress basal GTP turnover of G_s_ (Fig. 1A) suggests that RAMP2 does not fully prevent GCGR association with G_s_ but leads to unproductive coupling (Fig. S1D), which has been observed for other GPCRs^28^. To understand how RAMP2 induces this unproductive coupling with G_s_, we obtained a structure of the GCGR/RAMP2/G_s_ complex by cryoEM. In the smFRET experiments with RAMP2, performed at low concentrations of receptor and G_s_ for a short time (~30 minutes), RAMP2 binding resulted in nearly full inhibition of active state(s), though G_s_ is still able to populate the fully outward conformation to some extent (Fig. 2E, salmon) relative to the distribution observed for agonist-bound receptor in the presence of RAMP2 (Fig. S5B, purple). However, we can force GCGR/RAMP2/G_s_ complex formation by incubating at high concentrations for longer times with excess stabilizing Nb35 (Fig. S6A-C). We ensured any GCGR/G_s_ present in the sample was complexed with RAMP2 by pulling down directly on FLAG-tagged RAMP2. Co-complex formation was confirmed by size-exclusion chromatography (SEC) and SDS-PAGE (Fig. S6B-C). This GCGR/RAMP2/G_s_/Nb35 complex was subjected to cryoEM imaging to yield a final density map at a global nominal resolution of 2.90 Å (Fig. 3A, Fig. S6D, Table S1). We observed clear density for the agonist peptide and G_s_ but no density was obtained for RAMP2 (Fig. S6D). Additionally, though most parts of the receptor can be modeled unambiguously (Fig. S6E), some regions of GCGR cannot be built due to lack of density. These regions include the ECD, ICL3 and H8. While the GCGR ECD in the absence of RAMP2 exhibits well resolved density (Fig. 3C), in the presence of RAMP2 the GCGR ECD is not well-defined (Fig. 3B), consistent with experiments presented above showing RAMP2 enhances dynamics of this domain (Fig. 2B). The ECD of GCGR plays an important role in determining peptide potency^20^; the observed flexibility induced by RAMP2 is consistent with the decreased glucagon potency observed in cells (Fig. 1C).

**Fig 3.**
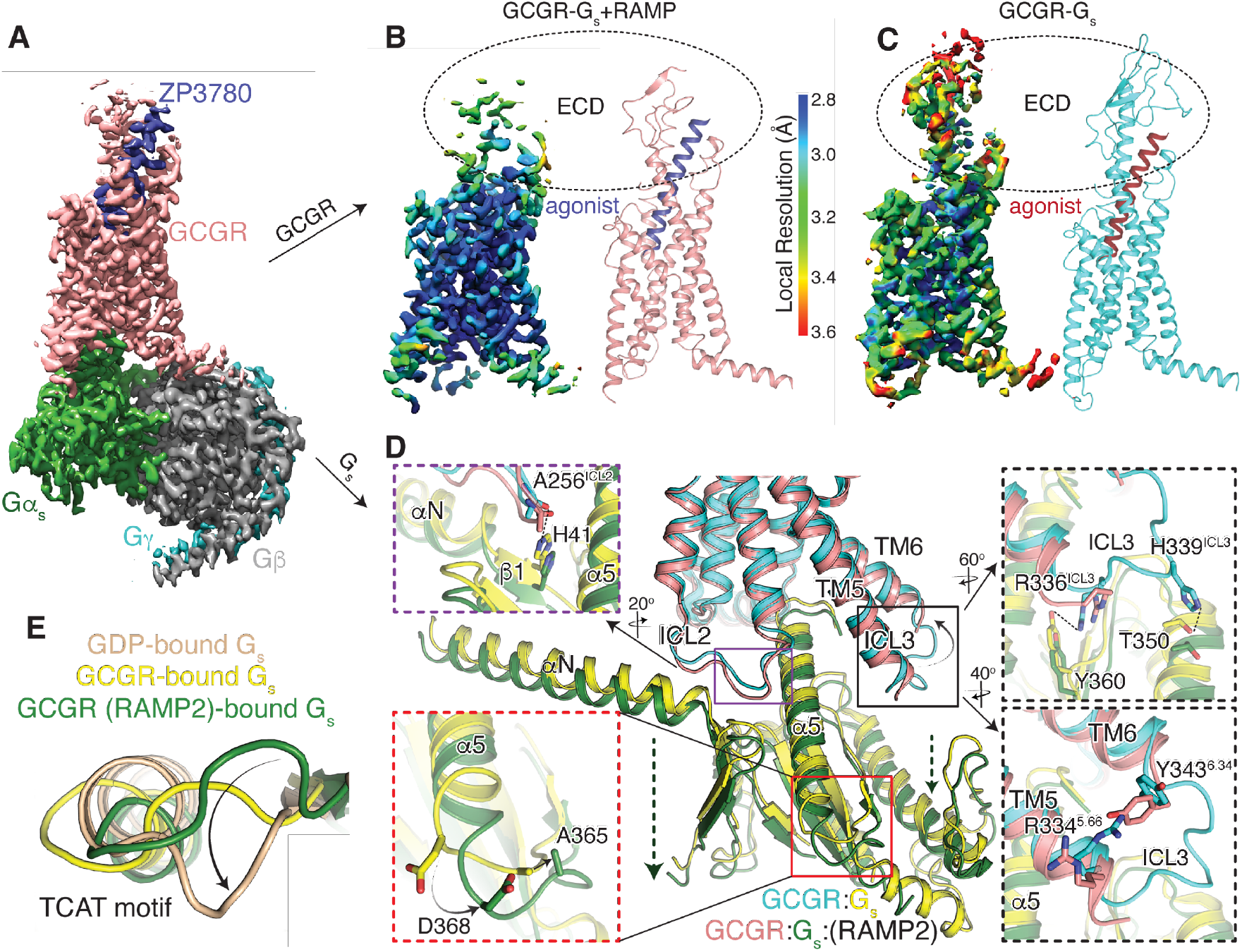
Structure of GCGR-G_s_ in the presence of RAMP2. (**A**) Cryo-EM density map of the ZP3780 and G_s_ heterotrimer bound GCGR in the presence of RAMP2 colored by subunit. Salmon, GCGR; blue, ZP3780; Gα_s_, green; Gβ, grey; Gγ, cyan. Cryo-EM density for ZP3780-bound GCGR in the presence (**B**) and absence (**C**) of RAMP2 colored by local resolution demonstrates a significant disordering of the receptor ECD in the presence of RAMP2. The intracellular interface of GCGR with Gα_s_ is perturbed by RAMP2 in several significant ways (**D**), including inducing disorder in ICL3 and the resulting loss of contacts with Gα_s_ (black, top), a downward movement of ICL2 (purple), and a lack of stabilizing contacts across TM5/TM6 (black, bottom). Further, there are rearrangements in the backbone and side chains in the TCAT motif at the base of the α5 helix of Gα_s_. Comparing this motif to GDP-bound G_s_ (**E**) it is clear that this important loop in the G protein occupies a distinct conformation.

Several specific interactions stabilizing the GCGR ECD are missing or modulated in the presence of RAMP2. A key disulfide bond between the N-terminal helix of the ECD (C43^ECD^) and a loop (C67^ECD^) is no longer observed and might be broken (Fig. S7A). This disulfide is highly conserved among Family B GPCRs and might be important to preserve the ECD fold^29^. Even regions that remain structured in the extracellular portion of the receptor are perturbed; a hydrogen bond between K37^ECD^ of the N-terminal ECD helix and S213^ECL1^ in ECL1 of the receptor is broken, along with a cation-*π* interaction stabilizing the N-terminal ECD helix (K35^ECD^/F31^ECD^) (Fig. S7B). The breaking of ECD interactions and specific structural elements affects peptide agonist/receptor interactions. E20 of the peptide agonist is no longer within hydrogen-bonding distance of Q131^1.29^ in TM1 (Fig. S7C), and deeper in the core of the receptor D195^2.68^ in TM2 forms an intrahelical salt bridge with R199^2.72^ at the expense of interaction with agonist, (Fig. S7D). The cryoEM density as well as the solution HDX-MS studies, which demonstrates increased ECD conformational sampling, suggest a linkage between perturbations in ECD conformation and intracellular TM6 conformation as observed by smFRET measurements; breakage and rearrangement of these interactions in the receptor core may provide a route for this negative allosteric communication.

ICL2 and ICL3 form an important part of the receptor-G protein binding interface (Fig. 3D)^3,30^. In the absence of RAMP2, ICL2 points into a hydrophobic pocket made up of residues in the αN/β1 hinge of Gα_s_ e.g. the carbonyl oxygen of A256^ICL2^ hydrogen bonds with H41 of Gα_s_ (Fig. 3D, purple box). However, in the presence of RAMP2, A256^ICL2^ is not within bonding distance of H41 of Gα_s_ (Fig. 3D, purple box), due to the entire G_s_ being ~ 3 Å further removed from the core of the receptor (Fig. 3D). Additionally, in the presence of RAMP2, no density is seen for ICL3 presumably due to its increased flexibility, which might be a result of losing stabilizing interactions across TM5-6 like the loss of a cation-π interaction between R334^5.66^ and Y343^6.34^ in the presence of RAMP2 (Fig. 3D, bottom black box). This ICL3 disorder results in the loss of a number of interactions with Gα_s_ such as H339^ICL3^ with T350 (Gα_s_) and R336 ^ICL3^ with Y360 (Gα_s_) (Fig. 3D, top black box). The lack of interaction with ICL3 could contribute to the observed G protein shift away from the receptor core.

Binding of G_s_ begins with the translation of α5 to engage receptor core. This leads to the rearrangement of the β6-α5 loop that contains the TCAT motif, which directly contacts the guanosine base of GDP^31^. The TCAT motif in G_s_ bound to GCGR in the presence of RAMP2 has moved ~5 Å (Cα of V365) away from the nucleotide-binding site compared to in the absence of RAMP2 (Fig. 3D, red box). This conformation of the TCAT might be more consistent with a conformation that cannot coordinate nucleotide. G_s_ bound to RAMP2-saturated GCGR displays a distinct TCAT motif relative to both GDP-bound G_s_ alone as well as GCGR-bound G_s_ (Fig. 3E). It is tempting to speculate that, in the presence of RAMP2, G_s_ adopts a conformation that cannot bind GTP and cause productive turnover, consistent with the G_s_-sequestering effect observed previously (Fig. S1D).

In order to better interpret the lack of local density in portions of the cryoEM structure of GCGR/G_s_ in the presence of RAMP2 we mapped the conformational heterogeneity in our final set of particles onto 3 principal components using 3D variability analysis (Fig. S8A)^32^. Within the principal components we see the typical “normal” motional modes observed for large biomolecular complexes (Fig. S8A; PC1, PC2). However, within the major principal component (PC0) there is evidence of concerted structural changes between extracellular and intracellular regions of the receptor. At one end of the continuum of structural snapshots (cyan), the density throughout the receptor and peptide is very comparable to the structure of GCGR/G_s_ in the absence of RAMP2 (Fig. 4A), including the ECD conformation “capping” the peptide agonist (red) and ICL3/H8 conformations consistent with fully G_s_-engaged, nucleotide-free complex^3^. At the alternate end of the PC0 continuum (salmon), the receptor ECD is in a completely distinct conformation having shifted away from the canonical peptide-bound conformation along with the top half of the agonist (blue). The TM region of the receptor does not display such conformational heterogeneity, consistent with the high resolution in the TM region (Fig. 3B). However, concurrent with the ECD movement away from the peptide binding region of the TM core, we observe H8 movement away from G_s_ upon loss of stable contacts (Fig. 4A). Such H8 contacts are important and common for Class B GPCR activity^33^. ICL3 has been shown to be important for receptor efficacy and G-protein selectivity^30,33^; it also occupies a distinct orientation in the alternate conformation observed from the principal component analysis. Thus, the likely explanation for the lack of structured density in the receptor ECD, ICL3, and H8 in our RAMP2-engaged complex is the presence of an ensemble of conformations in the final sample, ranging from those similar to the fully-active, canonical G_s_-bound state to a new, RAMP2-induced conformation lacking many contacts necessary for Class B GPCR signaling through G-proteins.

**Fig 4.**
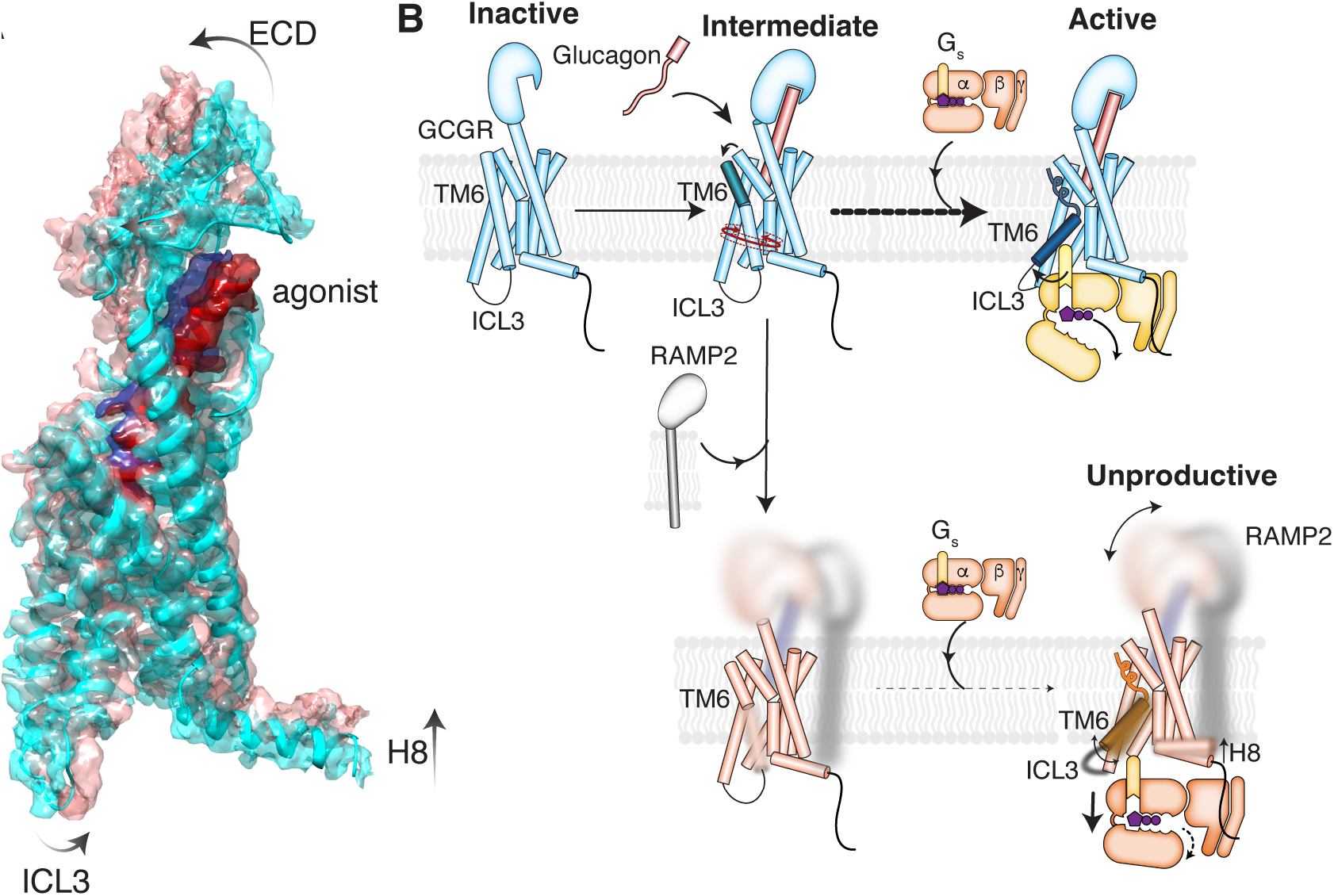
RAMP2-induced conformational dynamics in the GCGR ECD inhibits intracellular activation. (**A**) The major principal component of the 3D variability analysis directly demonstrates that changes in the GCGR ECD and agonist conformation are linked to changes in the intracellular conformation including ICL3 and H8. (**B**) Proposed model for RAMP2-induced inhibition of GCGR activation. Glucagon binding to GCGR promotes population of a newly characterized intermediate conformation of TM6, followed by full outward movement upon binding to G_s_. RAMP2 binding to GCGR causes enhanced ECD conformational sampling as observed in both HDX-MS solution experiments as well as cryo-EM structural analysis. This increase in dynamics in the ECD inhibits formation of both intermediate and active states of TM6 observed by smFRET experiments, and any engagement with G_s_ results in a largely unproductive complex.

## Discussion

In this work we present data on signaling modulation of a Family B GPCR, GCGR, by RAMP2. Unlike previous studies that have focused on obligate heterodimers forming RAMP-Family B GPCRs (e.g. CLR), our study investigates the effect of non-obligate heterodimer of RAMP2 and GCGR. We demonstrate using in-cell, biochemical and biophysical assays that RAMP2 interacts with GCGR and this interaction has profound implications for GCGR signaling. In agreement with previous studies in cells, we found that GCGR overexpression enhances cell surface levels of RAMP2^34^ and the presence of RAMP2 decreases glucagon potency at GCGR^11^. In a GTP-turnover assay, RAMP2 decreases GCGR-induced G_s_ activation in a GDP-dependent manner by slowing GDP release from the G protein. In addition to inhibiting G_s_, RAMP2 has been shown to decrease β-arrestin recruitment to GCGR^12^, and we present data to demonstrate that this is probably due to its inhibition of GRK5 phosphorylation. We show that RAMP2 broadly increases GCGR dynamics, including in the ECD, ICL3 and H8, and specifically stabilizes TM6 in an inactive state. Taken together our data suggests a model in which RAMP2 binding to GCGR increases ECD dynamics (Fig. 4B) which translates to decreased ligand potency. Several studies have shown single point mutations in the ECD, even if they are not in the direct ligand interacting residues, change ligand efficacy^20^. The role of Class B GPCR ECD conformation(s) in relation to intracellular activation remains underexplored, though there are likely differences between evolutionarily distinct subclasses of receptors. These differences in ECD conformations found in distinct subclasses of Class B GPCRs might alter their mode of interaction with RAMPs. For example, all three RAMPs interact with the calcitonin subfamily of receptors by “clamping” the receptor ECD in a manner where the extreme N-terminal helix is near parallel to the bilayer (Fig. S9A-C). To explore the conformation of GCGR ECD in the presence of RAMP2, we prepared covalently crosslinked GCGR/RAMP2/G_s_ complex and were able to observe low-resolution density for GCGR/RAMP2 containing micelles. These 2D classes showed a similar orientation to that predicted from *AlphaFold* (Fig. S8B) with the RAMP and GCGR ECDs appearing upright next to each other. Therefore, in contrast to calcitonin receptors, the ECD of peptide-bound glucagon receptor subfamily members is in an upright orientation with an N-terminal helix that is perpendicular to the bilayer (Fig. S9D). The GCGR ECD is probably prevented from occupying a “clamped” calcitonin-like conformation due to the differences in the helical nature of their peptide ligands, wherein glucagon is completely helical and calcitonin peptides are not.

RAMP2 binding appears to increase the disorder in the ECD of the receptor resulting in an enhanced population of the inward, inactive state of TM6 (Fig. 4B), though this does not seem to preclude G protein binding. The presence of RAMP2 slows GCCR-induced GDP release from G_s_ and inhibits GEF activity to levels lower than intrinsic basal nucleotide exchange; both suggest that interaction of RAMP2 with GCGR results in unproductive coupling to G_s_. Hence, RAMP2 is a negative allosteric modulator of GCGR whose allosteric effects appears to be driven by increasing flexibility in parts of GCGR ECD that translates to unproductive G_s_ coupling and thereby, signaling. This kind of “dynamic allostery” has been proposed for other RAMP complexes^5,8^ and such flexibility driven allosteric effects play an underappreciated role in protein– protein interactions in GPCRs. The unproductive GCGR-G_s_ coupling in the presence of RAMP2 seems to sequester G_s_ away from other GPCRs. Though the exact effect(s) of this G_s_ sequestration is unclear, a potentially interesting consequence may be found in glucagon signaling from the liver versus the kidney. We show here that RAMP2:GCGR ratios are important for achieving the observed changes in glucagon potency (Fig. 1C); the RAMP2:GCGR expression ratio is very low in liver (0.08:1) suggesting minimal inhibition and sequestration occurs, though in kidney the ratio is much higher (1.71:1)^35^ where the function of the glucagon receptor is not to regulate blood sugar but to modulate excretion of certain ions^36^. Differential expression of interacting proteins can significantly diversify biological functions^37^. Thus, the expression profiles of GCGR and RAMP2 may be uniquely titrated as a strategy to finely calibrate the way in which individual tissues, even individual cells, respond to the same extracellular stimulus. Understanding the effect of these GPCR-effector tissue specific interactions may allow for exploiting such variation as a source of targeted GPCR signaling output selectivity in drug development.

## Acknowledgements

We thank S. Reedtz-Runge (Novo Nordisk A/S) for providing the ligand NNC0640. We thank Betsy White for providing pure G-proteins. We thank S. Shoemaker and N. Dall for assistance with HDX-MS analysis scripts. K.K. was supported by the American Diabetes Association (ADA) Postdoctoral Fellowship. E.S.O. was supported by the American Heart Association (AHA) Postdoctoral Fellowship. N.R.L. was supported by a Miller Postdoctoral Fellowship. The work is supported by NIH grant GM050945 to S.M.; S.M. and B.K.K. are Chan Zuckerberg Biohub investigators.

## Author Contributions

K.K., D.H., E.S.O., and B.K.K. conceived the project. K.K., E.S.O and B.K.K. wrote the manuscript with input from all authors. K.K. and E.S.O. purified proteins, performed the GTPase GLO assay, fluorescence experiments, phosphorylation assays, and ^3^H-GDP release assays. K.K. and E.S.O collected cryoEM data with help from E.M. K.K. and E.S.O. processed cryoEM data with help from H.W. E.S.O. performed the 3D variability analysis. K.K. optimized labeling scheme and performed cwEPR experiments with help from D.H. J.M.M. developed the antagonist peptide, and optimized and performed cell based in vitro assays together with I.T. N.R.L. performed and analyzed HDX-MS experiments under the supervision of S.M. H.W. generated AlphaFold models. C.H. performed and analyzed smFRET experiments under the supervision of E.Y.I.

## Supplementary Material for

### Supplemental Figures

**Fig. S1.**
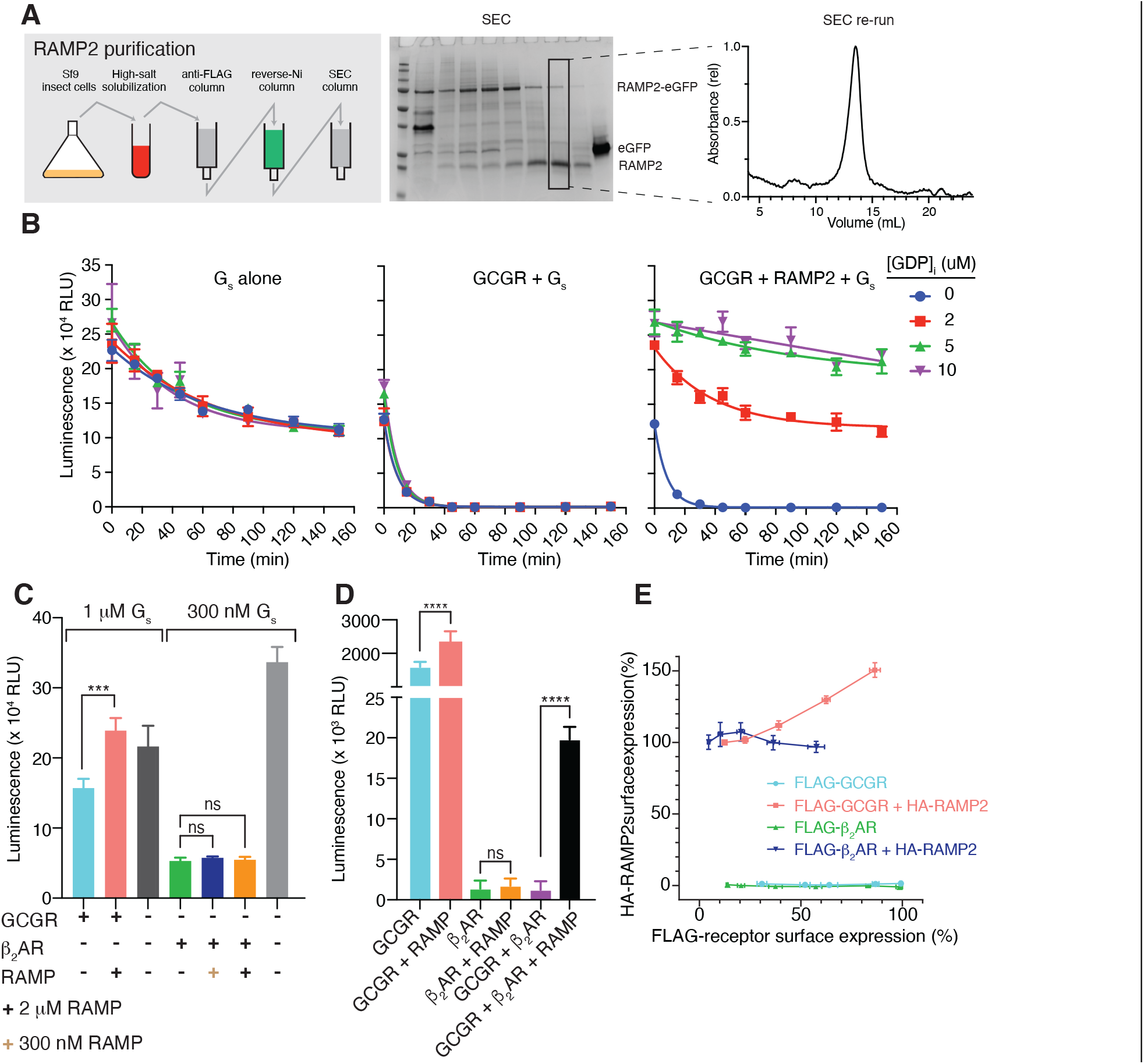
RAMP2 purification and effects on GCGR-induced G_s_ turnover. (**A**) Schematic of the RAMP2 growth and purification protocol, size exclusion chromatography profile demonstrating purity, as well as stability to repeated size exclusion runs. (**B**) Time-dependent GTP turnover assay demonstrating GDP-independent turnover for G_s_ alone and enhancement in turnover in the presence of agonist-bound GCGR. Addition of RAMP2 results in a [GDP] dependent inhibition in this turnover rate even past G_s_ alone. **(C)** GTP turnover assay demonstrating that RAMP2 inhibition is specific to GCGR and not observed for β_2_AR. **(D)** GTP turnover assay demonstrating that while RAMP2 has no direct effects on β_2_AR turnover, GCGR/RAMP2 complex is able to sequester away G_s_ from and inhibit β_2_AR dependent turnover. (**E**) Increasing cell-surface expression of glucagon receptor, but not β_2_AR, results in an increase in surface expression of RAMP2.

**Fig. S2.**
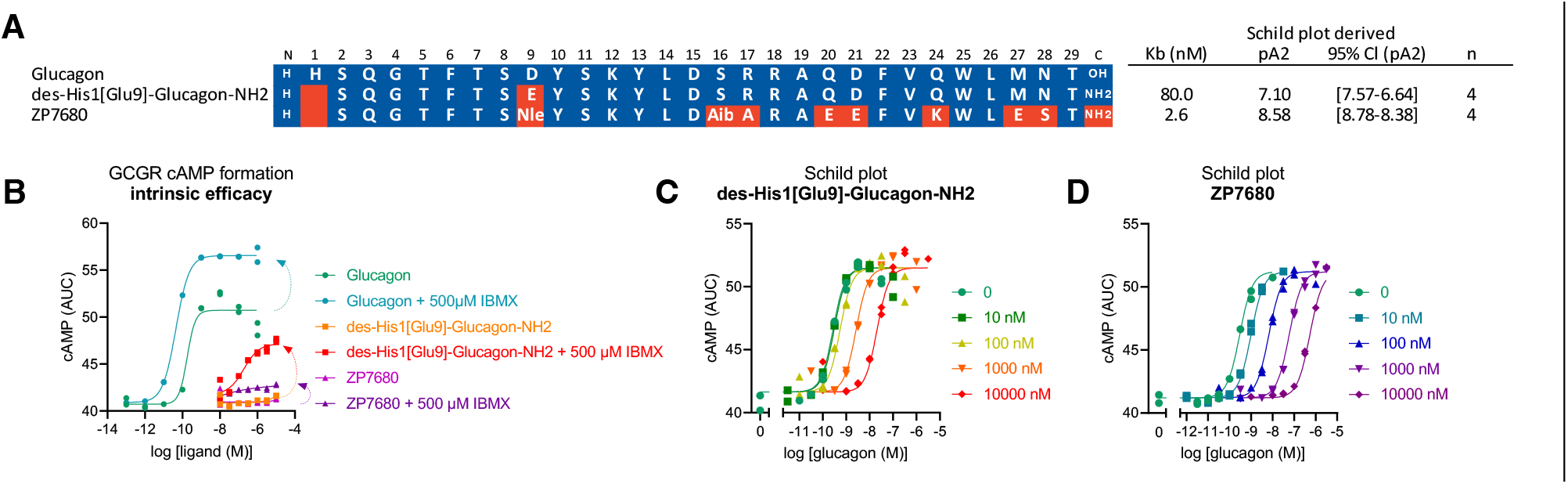
Design and characterization of the glucagon antagonist, ZP7680. (**A**) Sequences and estimated affinities of previously described GCGR antagonist des-His1[Glu9]-Glucagon-amide (K_B_ =80 nM), and new high affinity GCGR antagonist ZP7680 (K_B_ =2.6 nM). ZP7680 was obtained by removal of His1 and replacement of Asp9 with Nle and Ser11 with Ala to obtain a soluble and potent antagonist. (**B**) The antagonist ZP7680 has minimal intrinsic efficacy compared to des-His1[Glu9]-Glucagon-amide. Under conditions allowing for cAMP accumulation by addition of the PDE inhibitor IBMX, des-His1[Glu9]-Glucagon-amide have some agonistic activity whereas ZP7680 is silent. Schild plots were generated for the antagonist des-His1[Glu9]-Glucagon-amide (**C**) and ZP7680 (**D**) in absence of IBMX and indicated that ZP7680 is effective in blocking the glucagon response at lower concentrations compared to des-His1[Glu9]-Glucagon-amide, which is reflected in a higher estimated affinity (K_B_ / pA2) from a Schild plot analysis. Data in C and D are representative curves from one experiment performed in duplicate determinations from a total of 3 (C) or 4 (D) independent experiments, respectively.

**Fig. S3.**
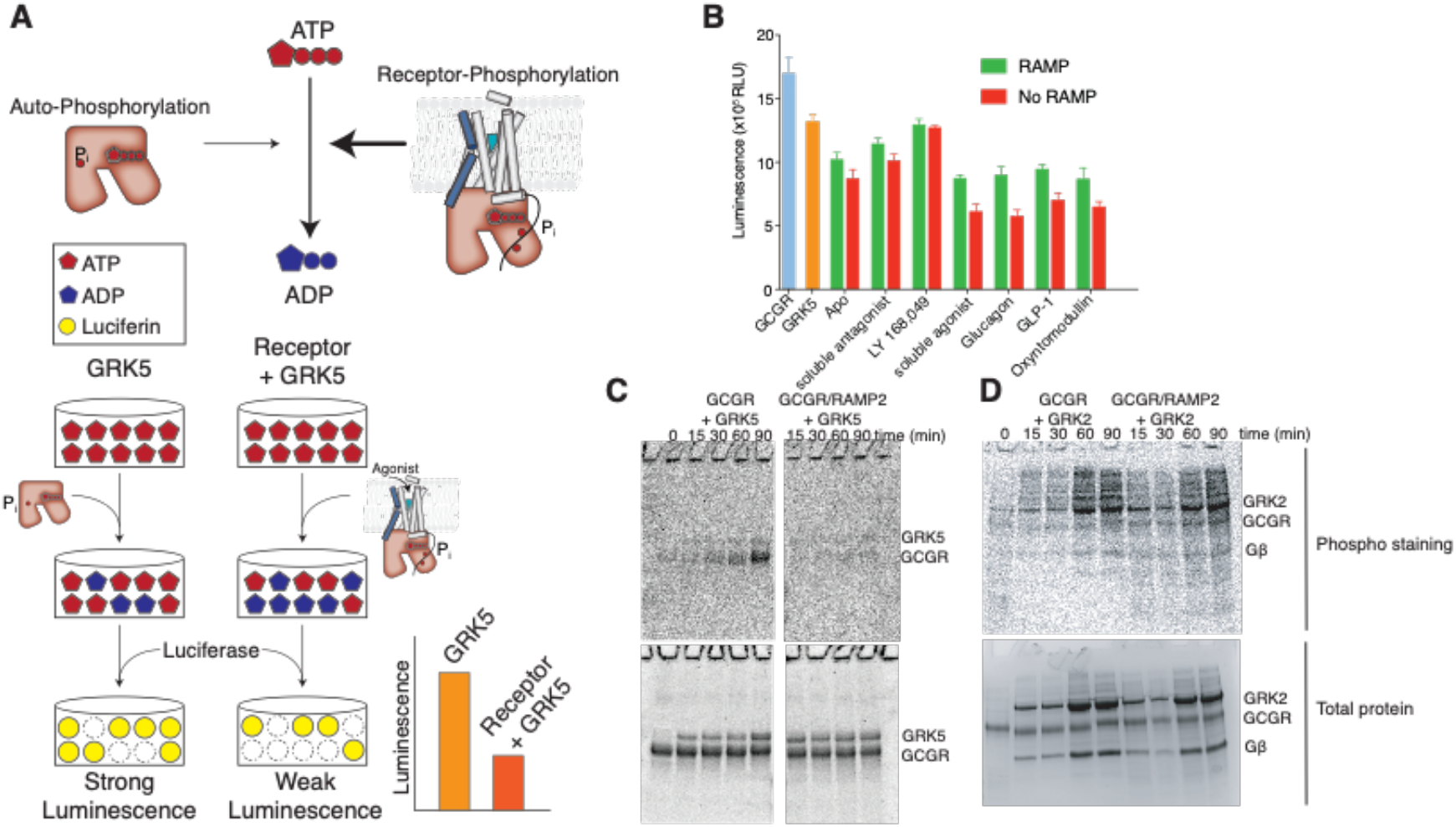
RAMP2 effects on GRK phosphorylation of GCGR. (**A**) Schematic depicting ATP depletion assay for kinase activity. (**B**) ATP depletion assay showing inhibition of GRK phosphorylation of GCGR in the presence of RAMP2. (**C**) Phosphorylation ProQ gels showing inhibition of GRK5 phosphorylation of GCGR by RAMP2 but (**D**) no inhibition of GCGR phosphorylation by GRK2.

**Fig. S4.**
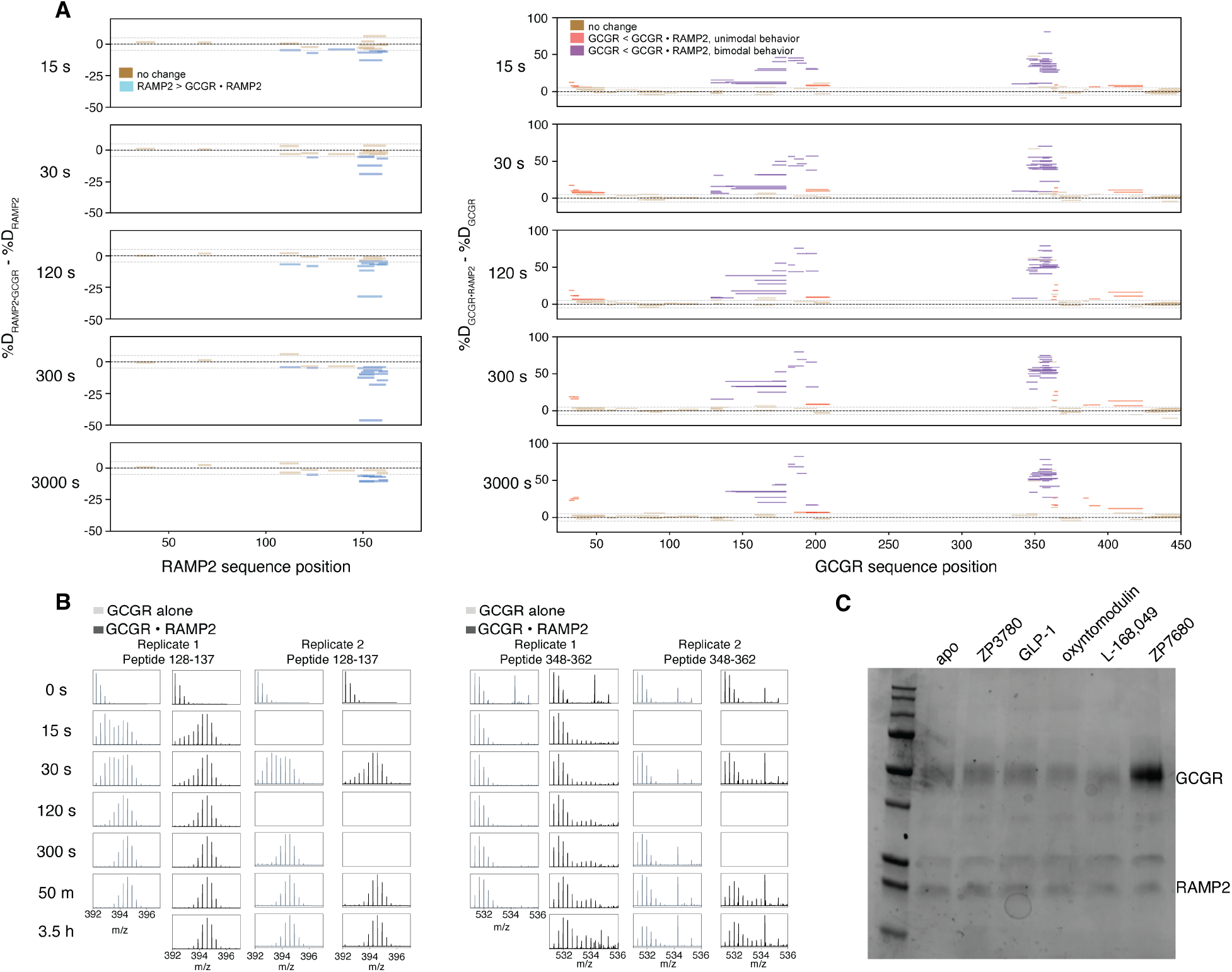
Perturbation in GCGR dynamics by RAMP2 observed by HX-MS. (**A**) Effect of binding partner addition on deuterium uptake for RAMP2 (+/- GCGR, left) and GCGR (+/- RAMP2, right). The difference in the percent deuteration for a given peptide at a given time point is plotted against the sequence position. Brown peptides show no difference in exchange, cyan peptides in RAMP2 are protected upon complexing with GCGR, and peptides that display unimodal and bimodal deprotection upon binding of RAMP2 are plotted in salmon and purple, respectively. (**B**) Replicate data for individual spectra of select peptides within regions of GCGR that display non-EX2 behavior as plotted in Fig. 2. (**C**) FLAG pulldown on RAMP2 assessed by SDS-PAGE gel demonstrating that antagonist peptide (ZP7680) bound GCGR interacts best with RAMP2.

**Fig. S5.**
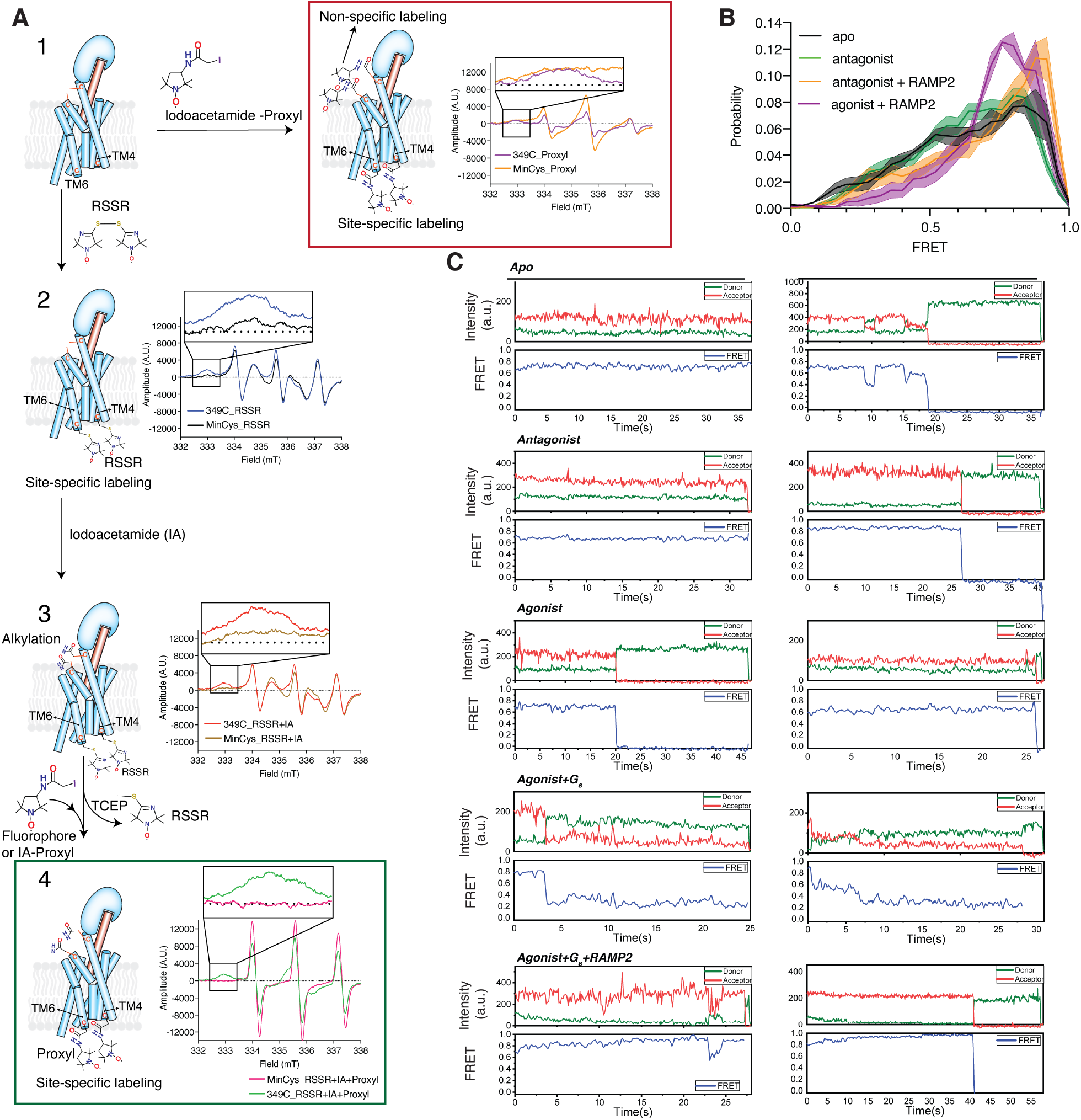
Site-specific labeling of mC-GCGR with cysteine-reactive groups and smFRET experiments. (**A**) Schematic highlighting the reaction scheme used to block labile, off-target disulfides and allow for specific labeling at introduced cysteines on GCGR. We use the highly cysteine-reactive nitroxide spin label Proxyl to test for off-target labeling by observing with cwEPR. Briefly, introduced cysteines are labeled specifically with the nitroxide spin label RSSR, followed by iodoacetamide capping of labile disulfides. Finally, RSSR is stripped off with TCEP, allowing for subsequent labeling with highly reactive fluorophores or spin labels of interest. (**B**) smFRET distributions of cysteine labeled GCGR comparing apo receptor to antagonist or antagonist and RAMP2 bound receptor, as well as the distribution observed for agonist and RAMP2 bound receptor. (**C**) Example smFRET traces showing donor (green), acceptor (red) intensity values as well as the calculated FRET values (blue) for a series of ligand conditions including apo, antagonist, agonist, agonist and G_s_, and agonist with G_s_ and RAMP2.

**Fig. S6.**
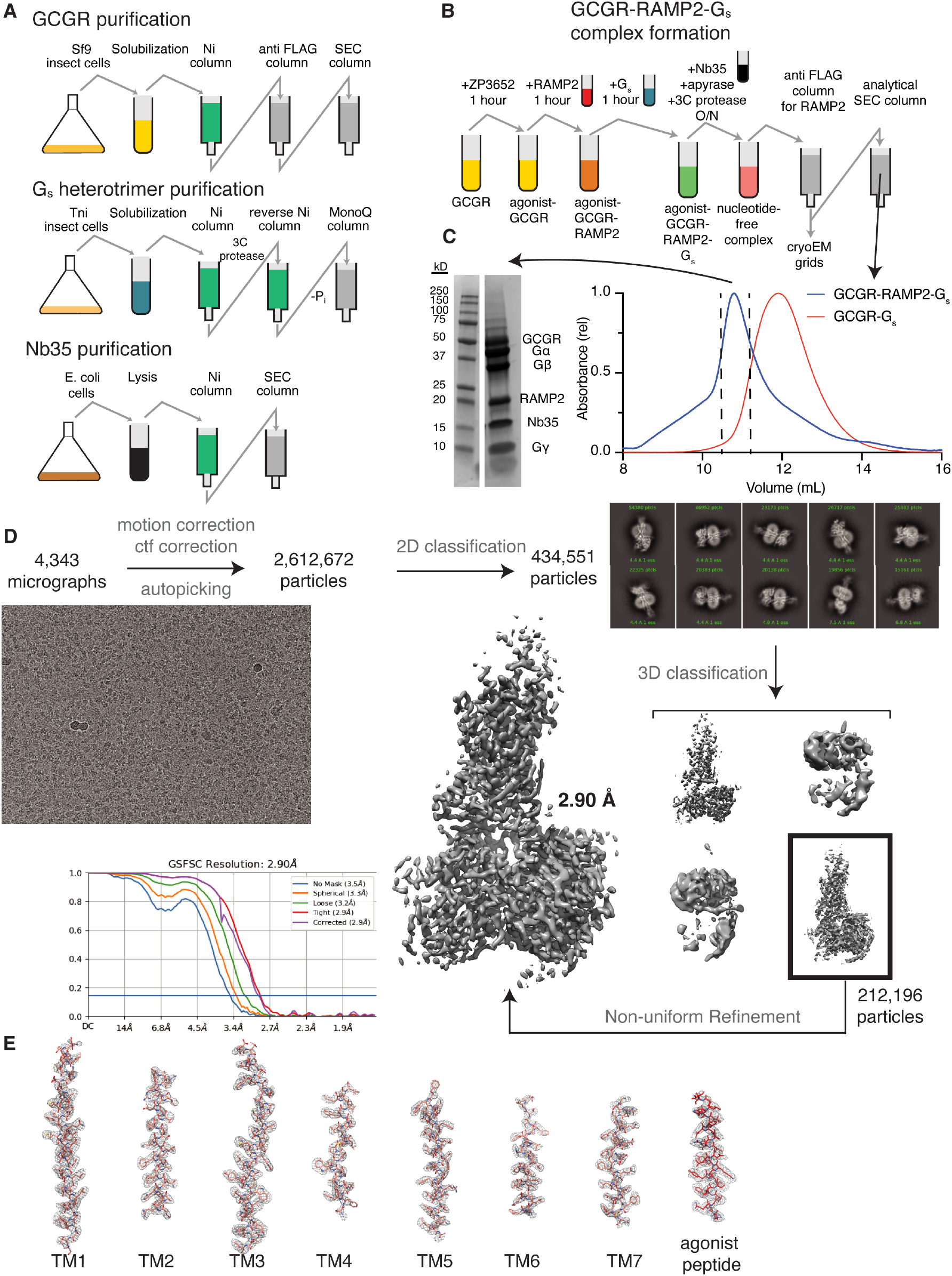
Cryo-EM sample preparation and data processing. Schematic depicting purification protocols (**A**) and complex formation (**B**). (**C**) Size exclusion chromatograms and SDS-PAGE gels demonstrating complex formation with RAMP2. (**D**) Cryo-EM data collection and processing pipeline, showing representative cryo-EM image of the GCGR-G_s_-RAMP2 complex, reference-free 2D cryo-EM averages, and cryo-EM data processing flow chart, including particle selection, classifications, density map reconstructions, and the “gold standard” FSC curves from cryoSPARC. (**E**) Cryo-EM density map and model are shown for the seven transmembrane helices and ZP3780.

**Fig. S7.**
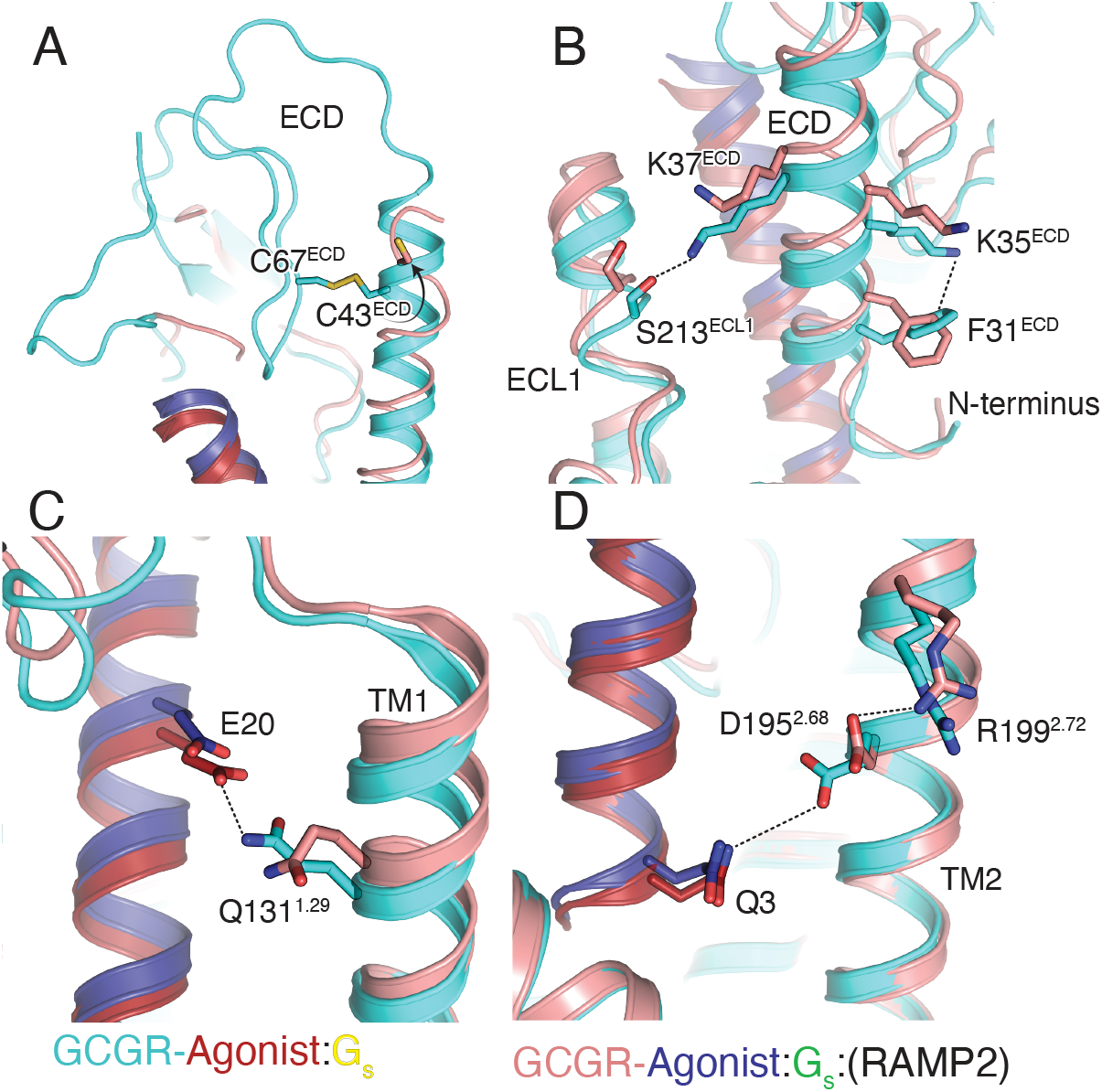
Broken ECD and agonist contacts in the presence of RAMP2. In the presence of RAMP2, there is no structured density for a conserved disulfide between C43 and C67 in the ECD (**A**), as well as a loss of a hydrogen bond between K37 in the ECD and S213 in ECL1 along with a cation-pi interaction within the N-terminal helix of the ECD (**B**). Ligand contacts with TM1 and TM2 are also lost, as seen in the loss of contact between E20 in the agonist and Q131 in TM1 (**C**) and D195 in TM2 switching hydrogen bond partners from Q3 in the agonist to R199 within TM2 (**D**).

**Fig. S8.**
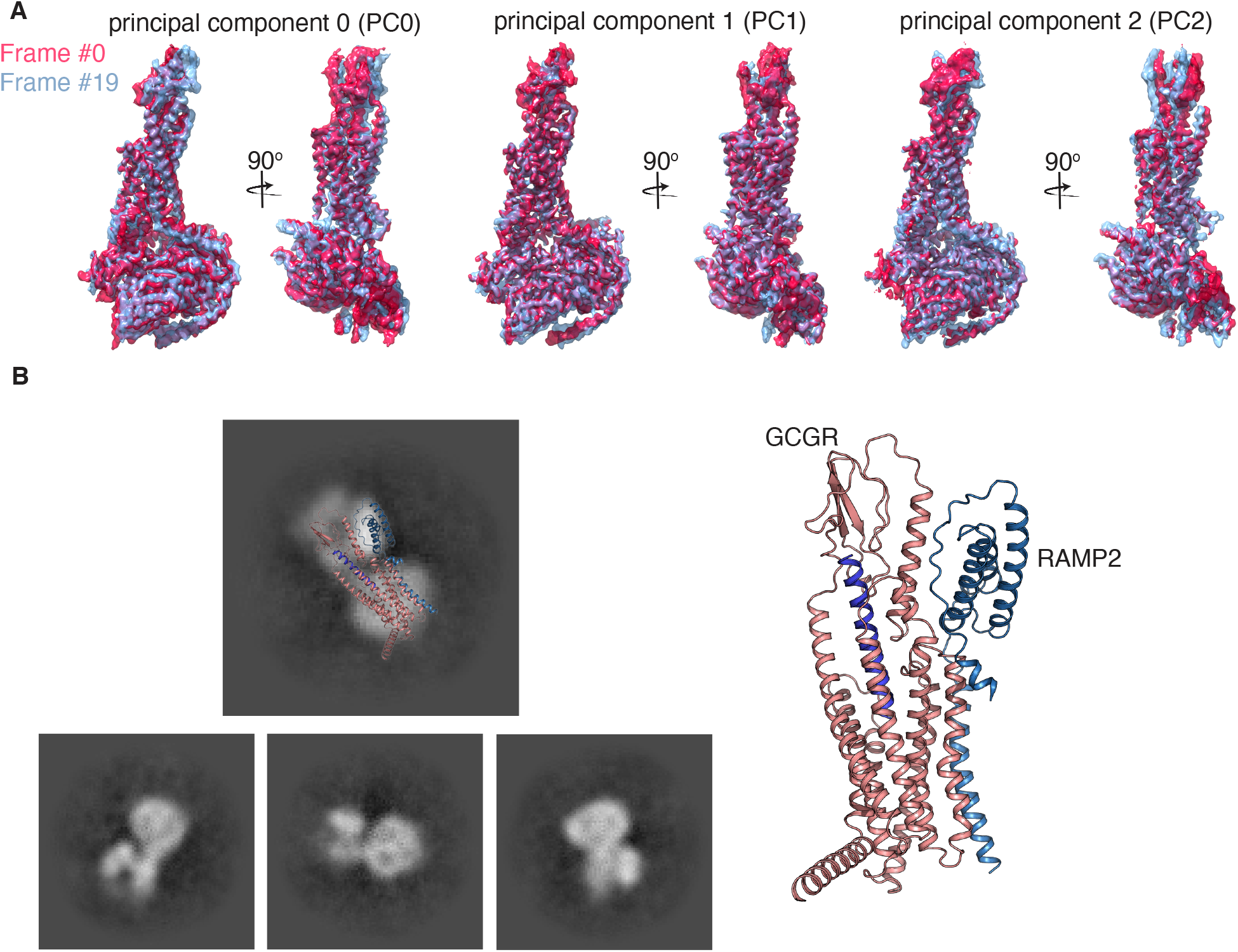
3D variability analysis and GCGR/RAMP2 ECD conformations. (**A**) The first (red) and last (light blue) frames within each of the 3 major principal components of the 3D variability analysis implemented in cryoSPARC. (**B**) Covalently crosslinked GCGR-RAMP2-G_s_ was subjected to cryo-EM imaging. The resulting particles were low-resolution, and only contained structured density for GCGR/RAMP2 or GCGR/G_s_ containing particles (not shown). The 2D reference-free class averages show density that corresponds well to the orientation predicted from AlphaFold for the GCGR/RAMP2 interface, i.e. both in an “upwards” conformation.

**Fig. S9.**
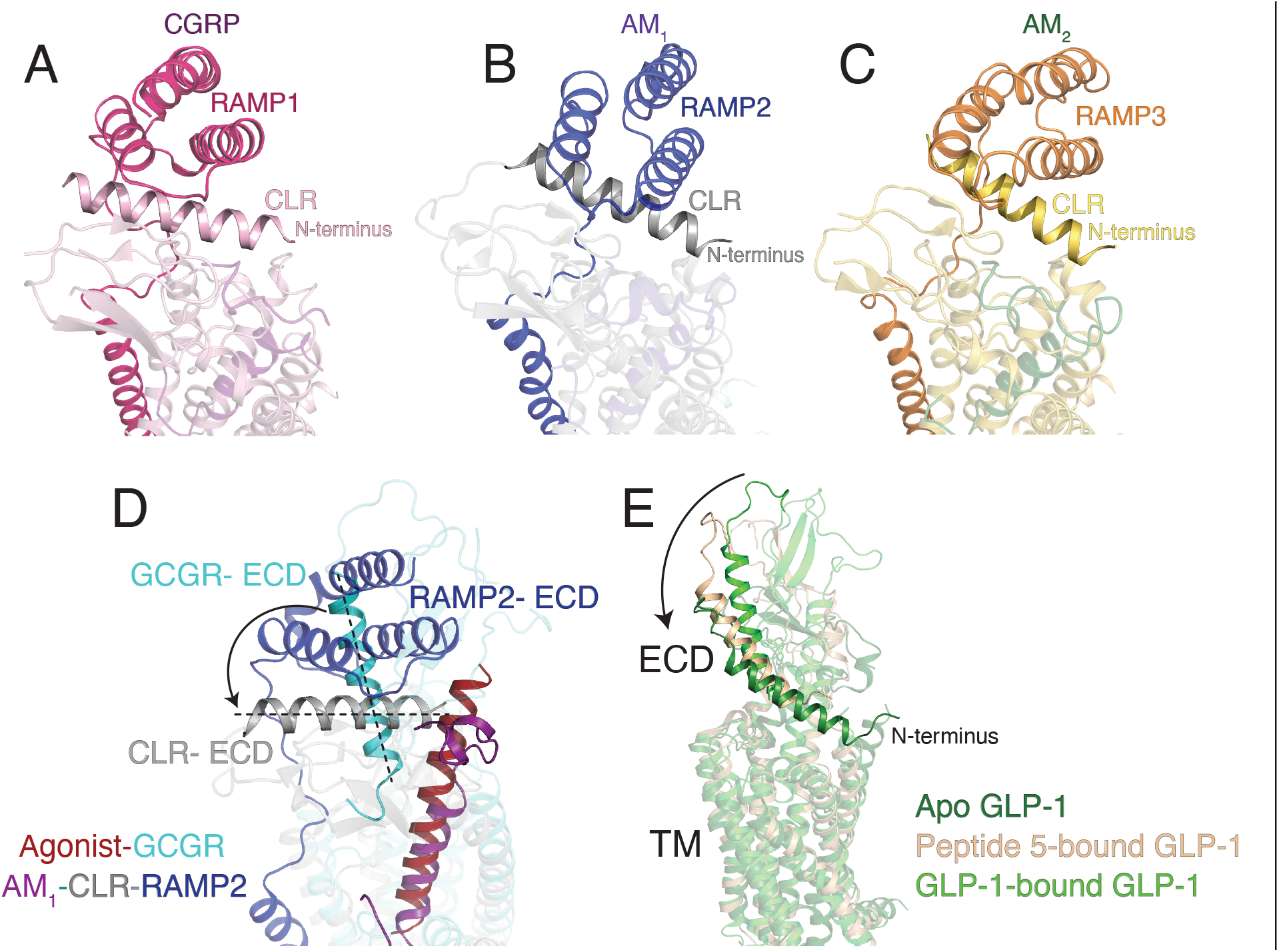
Differences in ECD conformation between glucagon-family and calcitonin-family Class B GPCRs. The structures of calcitonin-like receptor (CLR) bound to each of RAMP1 (CGRP, **A**), RAMP2 (AM1, **B**), or RAMP3 (AM2, **C**) display slight differences in binding pose but broadly bind to the CLR ECD with the N-terminal helix in an orientation near perpendicular to the bilayer norm. In contrast, the peptide-bound GCGR ECD is in an orientation that is incompatible with the same binding pose as that observed for CLR (**D**), even though there is some flexibility in this region in glucagon receptor family members (**E**).

### Supplemental Tables

**Table S1:** CryoEM data collection, model refinement and validation.

|  |  |
| --- | --- |
| **Data Collection** |  |
| Voltage (kV) | 300 |
| Magnification | 96,000x |
| Total electron dose (e<sup>-</sup>/Å<sup>2</sup>) | 56.6 |
| Defocus range (μm) | -0.7 to -2.0 |
| Calibrated pixel size (Å) | 0.852 |
| Micrographs collected | 4343 |
| **Data Processing** |  |
| Extracted particles | 2,154,997 |
| Particles used for final reconstruction | 212,196 |
| Final map resolution (Å, 0.143 FSC) | 2.90 |
| Map resolution range (Å) | 2.5 – 3.2 |
| Map sharpening B factor (Å<sup>2</sup>) | 103.2 |
| **Model Content** |  |
| Initial models used (PDB code) | 6WPW (GCGR/G<sub>s</sub>/Nb35) |
| Total number of atoms | 8,834 |
| No. of protein residues | 1,112 |
| No. of ligands | 0 |
| **Model Validation** |  |
| CC map vs. model (%) | 77 |
| RMSD |  |
| Bond lengths (Å) / Bond angles (°) | 0.002 / 0.598 |
| Ramachandran plot statistics |  |
| Favored (%) | 97.4 |
| Allowed (%) | 2.6 |
| Outliers (%) | 0.0 |
| Rotamer outliers (%) | 0 |
| C-beta deviations | 0 |
| Clash score | 3.83 |

### Materials & Methods

#### In vitro assay cell lines and constructs

HEK293 cells stably expressing an Epac1-based fluorescence resonance energy transfer (FRET) cAMP biosensor capable of reporting intracellular cAMP levels (cAMP biosensor cells) described previously^1^ were transfected with expression vector DNA encoding a FLAG-tagged human GCGR alone or together with HA-tagged RAMP2. The FLAG-tagged GCGR (FLAG-GCGR) consisted of a HA-signal peptide (MKTIIALSYIFCLVFA) followed by a FLAG tag (DYKDDDD), a small linker comprising a NotI restriction site (AAA) and the mature human GCGR (UNIPROT ID: P47871, residues 26-477). Likewise, a FLAG-tagged β_2_-adrenergic receptor was generated. The HA-tagged RAMP2 (HA-RAMP2) consisted of the HA-signal peptide followed by the HA-tag (YPYDVPDYA), a small linker comprising a NotI restriction site (AAA) and the mature human RAMP2 (UNIPROT ID: O60895, residues 36-175;). cAMP biosensor cells were maintained in growth medium (D-MEM, Gibco; supplemented with 10% fetal bovine serum, Gibco; 1% sodium pyruvate, Gibco; 1% MEM non-essential amino acids, Gibco; and 1% penicillin–streptomycin Solution, Gibco) and 50 ug/ml zeocin for selection of cell expressing the FRET-based cAMP biosensor construct.

For characterization of the antagonistic properties of ZP7680, the FLAG-tagged human GCGR was stably expressed in the HEK293 Epac1 biosensor cell line. A clone with an expression level corresponding to the transiently expressed FLAG-tagged human GCGR described above was used. The stable expressing cell line was maintained in growth medium with 50 ug/ml zeocin and 0.5 mg/ml G418, latter for selection of cells expressing the FLAG-GCGR construct.

#### cAMP assay

To assess the impact of RAMP2 on GCGR signaling, the cAMP biosensor cells were transiently transfected in suspension as described previously^2^ with different amounts of vector DNA encoding the FLAG-GCGR without or in presence of a fixed concentration of vector DNA encoding the HA-RAMP2. In brief, cAMP biosensor cells were brought into suspension by trypsination and resuspended in growth medium to a density of 200.000 cells/ml. For 1 ml of cell suspension transfected, 3 µL Fugene6 (Promega) in 57 µl OptiMEM (Gibco) was mixed with a total of 1 µg DNA in 25 µl OptiMEM. To vary surface expression levels (monitored by cell surface ELISA below) a range of 3-50 ng vector DNA encoding FLAG-GCGR supplemented with vector DNA to a total of 1 µg or with 800 ng vector DNA encoding HA-RAMP2 and vector DNA to a total of 1 µg was used. The Fugene6/DNA mixture was incubated for 20 min at room temperature and then added to 1 ml of cell suspension. Transfected cells were then seeded in black poly-D-lysine coated 96-well plates at 100 ul/well. Amounts were scaled up depending on the number of wells to be assayed. The cAMP assay was performed 48 hours after transfection. Prior to the assay, cells were washed once in 100 ul assay buffer (HBSS; Gibco, 20 mM HEPES pH 7.5, 1 mM CaCl_2_, 1 mM MgCl_2_, 0.1% BSA), and replaced with 100 µl of fresh assay buffer. After 15 min preincubation, cells were stimulated with glucagon diluted to different concentrations in 50 μl assay buffer and cAMP levels monitored continuously for 60 min by recording the change in FRET of the cAMP biosensor using an Envision plate reader (Perkin Elmer) equipped with filters for measuring the fluorescence of mCerulean fluorescent protein (474 nm) and mCitrine fluorescent protein (524 nm) following excitation of mCerulean (434 nm). To determine the EC_50_ and Emax of glucagon, the area under the curve for the individual cAMP traces was calculated and plotted against the concentration of glucagon and fitted by nonlinear regression to a 4-parameter logistic curve using GraphPad Prism.

#### Cell surface ELISA assay

To assess the expression level of GCGR and the impact of GCGR and RAMP2 on surface expression levels, the FLAG-tagged GCGR and HA-tagged RAMP2 constructs described above were used in a direct cell ELISA assay as described previously^3^. In brief, cells transfected for the cAMP assay above were seeded in white poly-D-lysine coated 96-well plates at 100 ul/well. At the same time as the functional assay, cells were fixated with 4% paraformaldehyde in DPBS, blocked in 3% dry milk in DPBS, incubated for 1 hour with HRP-conjugated anti-Flag antibody (Sigma Aldrich), or HRP-conjugated anti-HA antibody (R&D Systems), both diluted 1:2000 in 3% dry milk, followed by washing 4 time first in 3% dry milk in DPBS and then in DPBS. Surface expressed proteins were then quantified by addition of 60 μl DPBS buffer and 20 μl HRP substrate (Bio-Rad) per well, incubation for 10 min and detection of luminescence using an EnVision plate reader (Perkin Elmer). FLAG-GCGR luminescence was normalized to that of 50 ng vector DNA transfected and HA-RAMP2 luminescence normalized to that of 800 ng HA-RAMP2 vector DNA alone and plotted against each other.

#### In vitro characterization of the glucagon antagonist ZP7680

The glucagon antagonist ZP7680 was designed by removing His1 and replacing Asp9 with Nle and Ser11 with Ala^4^ in the dasiglucagon backbone^5^ to obtain a soluble and potent glucagon antagonist with minimal intrinsic efficacy. ZP7680 was characterized in vitro using the HEK293 Epac1 biosensor cell line stably transfected with the FLAG-tagged GCGR. Measurement of cAMP levels was determined as described above. Concentration response curves were generated in absence (as above) or presence of 500 µM IBMX to assess the intrinsic efficacy of ZP7680 and the previously described glucagon antagonist des-His1[Glu9]-Glucagon-NH2^6^. For generation of Schild plots to calculate antagonist K_B_ values, the antagonists were preincubated at different concentrations for 15 min prior to adding glucagon at increasing concentrations. No IBMX was added. The GraphPad Prism Gaddum/Schild EC50 shift function was used to determine antagonist K_B_-values.

#### Purification of wild type GCGR

Human glucagon receptor (Q27-F477) with an N-terminal FLAG and C-terminal octahistidine tag with 3C sites to remove tags was expressed as previously described^7^ using the baculovirus method in Spodoptera frugiperda (Sf9) cells. L-168,049 was added to 10 µM final upon infection and cells were collected 48 hours later and stored at −80°C until purification. GCGR was extracted from cell membranes with 1% lauryl maltose neopentyl glycol (L-MNG; Anatrace) and 0.1% cholesterol hemisuccinate (CHS) in 20 mM HEPES pH 7.5, 150 mM sodium chloride (NaCl), 20% glycerol, 5 mM imidazole, 30 µM NNC0640 ligand, and the protease inhibitors benzamidine and leupeptin, along with benzonase (Sigma-Aldrich). After douncing and stirring at 4 °C 1.5 hours, the solubilization mixture was centrifuged to pellet cell debris, and the supernatant was applied to nickel-chelating sepharose resin. After binding to nickel resin for 2 hours, the resin was washed with 20 mM HEPES pH 7.5, 150 mM NaCl, 0.1% L-MNG, 0.01% CHS, 20% glycerol, 5 mM imidazole, and leupeptin/benzamidine, followed by elution in the same buffer supplemented with 200 mM imidazole. 2 mM calcium chloride was added to the elution from the nickel resin and subsequently applied to M1 anti-FLAG immunoaffinity resin. The protein was washed in 20 mM HEPES pH 7.5, 100 mM NaCl, 0.05% L-MNG, 0.005% CHS with 2 mM calcium chloride, and eluted in the same buffer (without calcium chloride) with 5 mM ethylenediaminetetraacetic acid (EDTA) and FLAG peptide. Finally, monomeric GCGR was separated from multimers and aggregates via size exclusion chromatography on an S200 10/300 Increase gel filtration column (GE Healthcare) in 20 mM HEPES pH 7.5, 100 mM NaCl, 0.02% L-MNG and 0.002% CHS. Pure, monomeric GCGR was then spin concentrated to ~300 µM and flash frozen at −80 °C until further use.

#### Purification of minimal cysteine GCGR

The minimal cysteine version of GCGR (C171T, C240A, C287A, C401V) and related constructs used for bulk fluorescence and single-molecule FRET experiments (F31C, F245C introductions) were expressed as previously described^7^ in Expi293F cells. Constructs were transfected into Expi293F (Thermo Fisher) cells expressing the tetracycline repressor with the Expifectamine transfection kit (Thermo Fisher) following manufacturer’s directions with induction 2 days post-transfection with doxycycline (4 ug/mL and 5 mM sodium butyrate) with 1 µM L-168,049 (Tocris Bioscience). Pellets were frozen and stored at −80°C for later purification. Cells were dounced and solubilized with 20 mM HEPES pH 7.5, 100 mM NaCl, 20% glycerol, 1% L-MNG, 0.1% CHS, protease inhibitors and benzonase, followed by purification on anti-FLAG immunoaffinity chromatography. Following FLAG purification, minimal cysteine constructs were labeled with fluorophores as described below followed by size exclusion chromatography on Superdex 200 Increase 10/300 gel filtration column (GE Healthcare) in 20 mM HEPES pH 7.5, 100 mM NaCl, 0.02% L-MNG and 0.002% CHS. Aliquots of labeled protein were flash frozen and used as below.

#### Purification of RAMP2

Human RAMP2 (UNIPROT ID: O60895, residues 43-175) with an N-terminal FLAG tag, C-terminal eGFP followed by a hexahistidine tag along with a 3C protease site between RAMP2 and eGFP was cloned into the pFastBac vector. Protein was expressed using the baculovirus method in Sf9 cells as for GCGR. Pellets were frozen at −80°C until purification. Cells were solubilized with 20 mM HEPES pH 7.5, 750 mM sodium chloride, 20% glycerol, 1% *n*-dodecyl-ß-D-malopyranoside (DDM; Anatrace), protease inhibitors and benzonase and dounced on ice. After stirring at 4°C 1.5 hours, solubilized RAMP-eGFP was separated from cell debris by centrifugation. 2 mM Ca^2+^ was added to the supernatant before purification on M1 anti-FLAG immunoaffinity resin. Following FLAG purification, 3C protease was added 1:10 w/w with RAMP2-eGFP to cleave off the eGFP overnight at 4°C. Cleaved, monomeric RAMP2 was separated from eGFP and excess 3C via reverse nickel chromatography, followed by purification of monomeric RAMP2 with size exclusion chromatography on Superdex 200 Increase 10/300 gel filtration columns (GE Healthcare) in 20 mM HEPES pH 7.5, 200 mM NaCl, and 0.05% DDM. Aliquots of monomeric RAMP2 were flash frozen for later experiments.

#### Purification of GRKs

Full length wild type human GRK5 and bovine GRK2 were cloned into pVL1392 vector with C-terminal hexa-histidine tags for baculovirus production. GRKs were expressed and purified as previously described^8,9^. In brief, Sf9 cells were infected with BestBac baculovirus at a density of 4.0 × 10^6^ cells per mL and collected 48 hours post-infection. Cells were pelleted and frozen for later purification. Cells were dounced and solubilized with 20 mM HEPES pH 7.5, 300 mM NaCl, 1 mM dithiothreitol (DTT; Sigma-Aldrich), 1 mM phenylmethylsulfonyl fluoride (PMSF; Sigma-Aldrich), 1 mM EDTA, 0.02% Triton X-100 (Sigma), protease inhibitors and benzonase. Following 1.5 hour incubation at 4 °C, cell debris was pelleted by centrifugation. The supernatant was incubated with nickel-chelating sepharose resin for 2 hours at 4 °C after adding 20 mM imidazole and 5 mM magnesium chloride. Following washing nickel resin with 20 mM HEPES pH 7.5, 300 mM NaCl, 1 mM DTT, 1 mM PMSF, protease inhibitors and 0.05% DDM with 40 mM imidazole, the protein was eluted with the same buffer supplemented with 200 mM imidazole. The eluate was applied to a MonoS 10/100 column (GE Healthcare) for cation-exchange chromatography purification and eluted with a linear sodium chloride gradient. Monomeric kinase was purified by size exclusion chromatography on Superdex 200 Increase 10/300 gel filtration columns in 20 mM HEPES pH 7.5, 200 mM NaCl, 100 µM tris(2-carboxyethyl)phosphine) (TCEP; Sigma), and 0.05% DDM. Aliquots of purified human GRK5 and bovine GRK2 were flash frozen for later experiments.

#### Expression & purification of heterotrimeric G-proteins

Heterotrimeric G_s_ were expressed in *Trichoplusia ni (T. ni)* using the BestBac method (Expression Systems) and purified as previously described^7,10^. Briefly, two baculoviruses were used, one encoding the respective Gα subunit and the other encoding both Gβ_1_ and Gγ_2_, along with a histidine tag and HRV 3C protease site at the amino terminus of the β-subunit. *T. ni* cells were infected with the two baculoviruses for 48 hours and were subsequently harvested by centrifugation and lysed in 10 mM Tris, pH 7.5, 100 µM MgCl_2_, 5 mM β-mercaptoethanol (βME), 20 µM GDP and protease inhibitors. The resulting membranes were harvested by centrifugation and solubilized in a buffer containing 20 mM HEPES pH 7.5, 100 mM NaCl, 1% sodium cholate, 0.05% DDM, 5 mM MgCl_2_, 5 mM βME, 5 mM imidazole, 20 µM GDP and protease inhibitors. The solubilization reaction was homogenized with a Dounce homogenizer and incubated with stirring at 4 °C for 1.5 hours followed by centrifugation. The soluble fraction was loaded to Ni-chelated Sepharose and gradually detergent-exchanged into 0.1% DDM. The protein was eluted in the above buffer supplemented with 200 mM imidazole and dialyzed overnight into 20 mM HEPES pH 7.5, 100 mM NaCl, 0.1% DDM, 1 mM MgCl_2_, 5 mM βME and 20 µM GDP along with HRV 3C protease.

The 3C-cleaved heterotrimer was further purified away from 3C and uncleaved protein with reverse Ni chromatography and the resulting G protein was dephosphorylated with lambda protein phosphatase (NEB), calf intestinal phosphatase (NEB), and Antarctic phosphatase (NEB) in the presence of 1 mM manganese chloride (MnCl_2_). After dephosphorylation, the protein was bound to a MonoQ 10/100 GL column (GE Healthcare) in 20 mM HEPES pH 7.5, 50 mM NaCl, 1 mM MgCl_2_, 0.05% DDM, 100 µM TCEP, and 20 µM GDP and washed in the same buffer, followed by elution with a linear gradient from the loading buffer to the same buffer with 500 mM NaCl. The main peak with G protein heterotrimer was collected and dialyzed into 20 mM HEPES pH 7.5, 100 mM NaCl, 0.02% DDM, 100 µM TCEP and 20 µM GDP overnight, followed by spin-concentration to <250 µM, addition of 20% glycerol, and flash-freezing in liquid nitrogen and storage at −80°C until further use.

#### Purification of Gα_s_ and Gβ_1_γ_2_

Gα_s_ and Gβ_1_γ_2_ were expressed and purified for the GDP release assay as previously described^7^. Gβ_1_γ_2_ was expressed and purified in an identical manner to the heterotrimer purification above (without co-infection with Gα_s_ virus) aside from the linear elution gradient from MonoQ 10/100 GL column (GE Healthcare), which for Gβ_1_γ_2_ is from the loading concentration of 50 mM NaCl to a final 250 mM NaCl concentration over 7.5 column volumes. Human Gα_s_ subunit with an amino-terminal hexahistidine tag with a HRV 3C protease site to cleave off the histidine tag was expressed in Rosetta 2 (DE3) cells (EMD Millipore) in a pET28a vector. Cells transformed with the above vector were grown in Terrific Broth to an OD600 of ~0.6 followed by induction of protein production with 0.5 mM isopropyl B-D-1-thiogalactopyranoside (IPTG; Sigma-Aldrich). After growth overnight at room temperature, cells were harvested and resuspended in lysis buffer composed of 50 mM HEPES pH 7.5, 100 mM NaCl, 1 mM MgCl_2_, 50 µM GDP, 5 mM βME, 5 mM imidazole and protease inhibitors. Cells in lysis buffer with subjected to sonication with a 50% duty cycle, 70% power for 4 cycles of 45 seconds. Intact cells and cell debris was removed with centrifugation, and the resulting supernatant was bound to Ni-chelated Sepharose resin and washed several times with lysis buffer in batch, followed by several column volumes washing on-column and elution with lysis buffer supplemented with 200 mM imidazole. Ni-purified protein was dialyzed overnight at 4°C against 20 mM HEPES pH 7.5, 100 mM NaCl, 1 mM MgCl2, 20 µM GDP, 5 mM βME and 5 mM imidazole along with HRV 3C protease to cleave off the amino-terminal histidine tag. Cleaved Gα subunit was passed through Ni-chelated sepharose resin to remove uncleaved protein, 3C protease, and amino-terminal histidine tags. The protein was then concentrated and run on a Superdex 200 10/300 Increase column in 20 mM HEPES pH 7.5, 100 mM NaCl, 1 mM MgCl_2_, 20 µM GDP, and 100 µM TCEP.

#### Purification of Nb35

Nb35 was expressed & purified as previously described ^11^. Briefly, plasmid was transformed into *Escherichia coli* BL21 cells and grown to an optical density 0.7-1.0 and expression was induced with 1 mM IPTG overnight at room temperature. Cells were harvested and lysed followed by purification via nickel affinity chromatography and size exclusion chromatography on a Superdex S200 10/300 Increase gel filtration column (GE Healthcare) in 20 mM HEPES pH 7.5 with 150 mM sodium chloride. Purified Nb35 was flash frozen and stored at −80°C for later use.

#### Formation & purification of GCGR-RAMP2-G_s_-Nb35 complex

GCGR-RAMP2-G_s_-Nb35 complex was formed as previously described for GCGR-G_s_-Nb35^7^ complex aside from the addition of separately purified RAMP2 (described above). 1% L-MNG/0.1% CHS was added to purified Gα_s_ß_1_γ_2_ for 1 hour on ice in order to exchange detergents from the DDM used for initial purification. ZP3780 was dissolved to 5 mM in dH_2_O and added to purified apo-GCGR to 500 µM final and incubated at room temperature for 1 hour. A 1.2-fold molar excess of agonist-bound GCGR was added to purified RAMP2 and incubated for an additional hour at room temperature. Detergent-exchanged Gα_s_ß_1_γ_2_ was then added to the GCGR-ZP3780-RAMP2 complex at a 1.5-fold molar excess with receptor and incubated at room temperature for 2 hours before addition of a 2-fold molar excess of Nb35 (with respect to Gα_s_ß_1_γ_2_) and further incubation on ice for 1.5 hours. Finally, apyrase (1 unit, NEB) and 3C protease (1:10 w/w ratio with GCGR) were added to stabilize the nucleotide-free state of Gα_s_ß_1_γ_2_ and cleave off the M1 FLAG tag of the receptor. The final molar ratios of protein components was 1:1.2:1.8:3.6 (RAMP2:GCGR:G_s_:Nb35). The complex was purified with M1 anti-FLAG affinity chromatography as previously described^7^ but using the M1 FLAG tag on the RAMP2 protein rather than on GCGR to ensure that any GCGR/G_s_ complex present was bound to RAMP2. Briefly, a 4-fold volume of 20 mM HEPES pH 7.5, 100 mM sodium chloride, 0.8% L-MNG/0.08% CHS, 0.27% glyco-diosgenin (GDN; Anatrace)/0.027% CHS, 1 mM magnesium chloride, 10 µM ZP3780, and 2 mM calcium chloride was added to the complexing reaction, followed by purification with M1 anti-FLAG chromatography. The complex was loaded to M1 anti-FLAG resin equilibrated with 20 mM HEPES pH 7.5, 100 mM sodium chloride, 0.3% L-MNG/0.03% CHS, 0.1% GDN/0.01% CHS, 5 µM ZP3780, and 2 mM calcium chloride and washed with 2 column volumes of the same. The complex was further washed with 4 additional column volumes of the same buffer with progressively lower detergent concentrations, followed by elution with 20 mM HEPES pH 7.5, 100 mM sodium chloride, 0.00075% L-MNG/0.000075% CHS, 0.00025% GDN/0.000025% CHS, 10 µM ZP3780, 5 mM EDTA, and FLAG peptide. The eluted complex was supplemented with 100 µM TCEP, analyzed by gel and size exclusion chromatography and concentrated to ~9 mg ml^−1^ for electron microscopy.

#### Crosslinking of the GCGR/RAMP2/G_s_ complex

The complex above containing GCGR, agonist, RAMP2, G_s_, and Nb35 was covalently crosslinked by adding disuccinimidyl dibutyric urea (DSBU, Thermo Scientific) to a final concentration of 3 mM for 1 hour on ice. Covalent crosslinking was confirmed by SDS-PAGE gel analysis as observed by a loss of monomeric complex components, and the resulting crosslinked complex was analyzed by size exclusion chromatography to confirm a lack of multimeric or aggregated species (data not shown).

#### Cryo-EM data acquisition & processing

An aliquot of purified GCGR-RAMP2-G_s_-Nb35 complex was applied to glow-discharged 200 mesh grids (Quantifoil R1.2/1.3) at a concentration of ~9 mg ml^−1^ and vitrified using a Vitrobot Mark IV (Thermo Fisher Scientific) at 100% humidity at 4 °C after blotting for 3 s with a blot force of 3. CryoEM images were collected on a Titan Krios operated at 300 kV at a nominal magnification of 96,000x using a Gatan K3 Summit direct electron camera in counting mode corresponding to a pixel size of 0.85 Å. A total of 4343 image stacks were obtained with a dose rate of 16.4 e^−^/pixel/s and total exposure time of 2.5 s with 0.05 s per frame, resulting in a total dose of 56.6 electrons per Å^2^. The defocus range was set to −0.7 to −2.0 µm.

All subsequent processing was performed with CryoSPARC^12^. Dose-fractionated image stacks were imported to CryoSPARC and subjected to patch-based beam-induced motion correction and contrast transfer function (CTF) estimation. A total number of 2,612,672 particles were picked with a template-based auto-picking protocol, followed by removal of particles from micrographs with a CTF resolution estimation of greater than 3.5 Å. A subset of 434,551 particles were selected after 2 rounds of 2-dimensional (2D) classification. These particles were further subjected to 3-dimensional (3D) classification with heterogeneous classification into 4 classes, resulting in a class containing 212,196 final particles, which were then reconstructed to 2.90 Å nominal resolution at FSC of 0.143 with non-uniform refinement^13^. Local resolution was estimated within CryoSPARC followed by 3D-variability analysis^14^ of the final particle set into 3 motional modes filtered to 5 Å.

#### Phosphorylation ATP depletion assay

GCGR was diluted to 40 µM in phosphorylation buffer composed of 100 mM HEPES pH 7.5, 35 mM sodium chloride, 5 mM magnesium chloride, 0.01% L-MNG/0.001% CHS, 20 µM diC8-PtdIns(4,5)P_2_, 100 µM TCEP and 80 µM ATP. Various ligands were added to 150 µM and bound to receptor at room temperature for 1 hour. Aliquots were split in half and RAMP2 added 0.75:1 with receptor and incubated for a further 1 hour at room temperature. GRK5 was diluted to 4 µM in the phosphorylation buffer above without ATP and incubated on ice until the start of the reaction, when equal volumes of receptor solution with different ligands and RAMP2 and GRK5 solution were mixed. Total ATP was measured by mixing receptor with phosphorylation buffer in the absence of GRK5, and intrinsic GRK5 activity was measured by mixing GRK5 solution with phosphorylation buffer without receptor. The reaction was allowed to proceed for 90 minutes at room temperature before quenching with equal volume 20 mM HEPES pH 7.5, 400 mM NaCl, and 10 mM EDTA on ice. Residual ATP was measured by adding equal volume of GTPase-GLO™ luciferase detection reagent for 10 minutes, and luminescence detected on SpectraMax Paradigm plate reader. The method is described graphically in Fig. S2. Raw luminescence values were converted to percent depleted values by normalizing to receptor alone (0% depletion) and zero signal (100% depletion).

#### Phosphorylation gel assay

GCGR (2.0 µM final) was equilibrated in phosphorylation buffer (100 mM HEPES pH 7.5, 35 mM sodium chloride, 5 mM magnesium chloride, 0.01% L-MNG/0.001% CHS, 20 µM diC8-PtdIns(4,5)P_2_, 100 µM TCEP and 1 mM ATP) and agonist ZP3780 was added to 10 µM final. For samples with GRK2, purified Gß_1_γ_2_ was added to 1 µM final. RAMP2 was added to 5 µM final. The reaction was equilibrated for 1 hour at room temperature to allow binding of agonist and RAMP2 to receptor before kinase (GRK2 or GRK5) was added in 200 nM increments (1:10 with GCGR each addition). The reaction(s) were sampled at various time points by quenching with 4x Laemmli Sample Buffer (Bio-Rad) with βME. Samples were analyzed by gel electrophoresis and stained with Pro-Q™ Diamond phosphoprotein gel stain (Thermo Fisher). Total protein was measured by staining with Coommassie blue or SYPRO Ruby (Thermo Fisher) protein gel stain (Thermo Fisher). Pro-Q™- and SYPRO™-stained gels were imaged with a Typhoon laser-scanner (Cytiva) using a 532 nm excitation wavelength and a 560 nm longpass filter.

#### GTP turnover assay

The GTP turnover assay was done using a modified version of the GTPase-GLO™ assay (Promega) as previously described^7,10^. Briefly, the final reaction was composed of 20 mM HEPES pH 7.5, 100 mM sodium chloride, 10 mM magnesium chloride, 100 µM TCEP, 0.01% L-MNG/0.001% CHS, variable concentrations of GDP, 10 µM GTP. Prior to initiating the reaction, GCGR was incubated in the presence of excess (20 µM) agonist ZP3780 in buffer with 20 mM HEPES pH 7.5, 100 mM sodium chloride, 0.01% L-MNG/0.001% CHS and 20 µM GTP for 1 hour in order to bind agonist. Concurrently, G protein was exchanged to L-MNG/CHS by adding 1% L-MNG/0.1% CHS for 1 hour on ice followed by dilution to 2 µM into G-protein buffer containing 20 mM HEPES pH 7.5, 100 mM sodium chloride, 20 mM magnesium chloride, 200 µM TCEP, 0.01% L-MNG/0.001% CHS and various concentrations of GDP. The solutions containing G_s_ and GCGR at 2-fold their final reaction concentrations were mixed to initiate the reaction. After incubation for various times (Fig. S1B) GTPase-Glo reagent supplemented with 10 µM adenosine 5’-diphosphate (ADP) was added and incubated with the sample for 30 minutes at room temperature. Luminescence was measured after addition of detection reagent for 10 minutes at room temperature using a MicroBeta Counter or using a SpectraMax Paradigm plate reader.

#### GDP release assay Gα_s_ and Gβ_1_γ_2_

The single turnover GDP release assay was performed similar to previously described^7^ with slight modifications. Briefly, Gα_s_ in 20 mM HEPES pH 7.5, 100 mM sodium chloride, 1 mM magnesium chloride, 100 µM TCEP and 5 µM GDP was diluted to 0.8 µM in GDP-loading buffer (20 mM HEPES pH 7.5, 100 mM sodium chloride, 1 mM EDTA and 100 µM TCEP) with 2.5 µM [^3^H]-GDP (Perkin Elmer). Gα_s_ was incubated in GDP-loading buffer for 1 hour at room temperature, after which, purified Gß_1_γ_2_ was added to a 1.2-fold molar excess and allowed to form complex with Gα_s_ for 30 minutes at room temperature. GCGR was diluted to 10 µM in 20 mM HEPES pH 7.5, 100 mM sodium chloride, 0.01% MNG/0.001% CHS, and 2 mM GTP and incubated with 100 µM ZP3780 for 1 hour at room temperature. The reaction was initiated by mixing the agonist-bound GCGR with [^3^H]-GDP loaded G_s_ heterotrimer at final concentrations of 5 µM and 200 nM, respectively. The reaction was stopped at various time points by dilution of the 20 µL reactions with 500 µL ice-cold wash buffer (20 mM HEPES pH 7.5, 150 mM sodium chloride, 20 mM magnesium chloride) and immediately filtered using a microanalysis filter holder (EMD Millipore) and pre-wet mixed cellulose filters (25 mM, 0.22 um). The filter was subsequently washed three times with 500 µL ice-cold wash buffer. Radioactivity was measured by liquid scintillation spectrometry.

#### RAMP2/GCGR pulldown assay

GCGR was diluted to 15 µM in 20 mM HEPES pH 7.5, 100 mM sodium chloride and 0.01% MNG/0.001% CHS, 5 mM magnesium sulfate and 2 mM calcium chloride along with a 1:10 w/w amount of 3C protease to cleave off the FLAG tag on GCGR. RAMP2 was added to ~10 µM final. The sample was split and various ligands (full agonist ZP3780, partial agonists GLP-1 and oxyntomodulin, negative allosteric modulator L-168,049, and antagonist peptide ZP7680) were added to 150 µM (10-fold molar excess with GCGR) final concentration and incubated overnight on ice. The next day, samples were purified on M1 anti-FLAG affinity resin using the residual M1 FLAG tag on the RAMP2. Samples were bound to M1 anti-FLAG resin equilibrated with 20 mM HEPES pH 7.5, 100 mM sodium chloride and 0.01% MNG/0.001% CHS and 2 mM calcium chloride for 20 minutes at room temperature, followed by washing over 15 minutes with ~5 column volumes of the same buffer to remove unbound GCGR. Samples were eluted with 20 mM HEPES pH 7.5, 100 mM sodium chloride and 0.01% MNG/0.001% CHS, 5 mM EDTA and FLAG peptides and analyzed by SDS-page chromatography (Fig. S4A).

#### Bulk fluorescence experiments

##### Labeling with IANBD-amide

Purified minimal cysteine GCGR with introduced cysteines at either position 345 (F345C, TM6) or position 31 (F31C, ECD) were diluted to 50 µM in 20 mM HEPES pH 7.5, 100 mM sodium chloride and 0.02% MNG/0.002% CHS. N,N’-dimethyl-N-(iodoacetyl)-N’-(7-nitrobenz-2-oxa-1,3-diazol-4-yl)ethylenediamine (IANBD-amide; Thermo Fisher) was solubilized in DMSO to make a 25 mM stock, then added to GCGR at 5-fold molar excess (250 µM) and incubated at room temperature 45 minutes. The reaction was quenched with excess cysteine and the reaction mixture was purified by size-exclusion chromatography on a Superdex 200 Increase 10/300 gel filtration column in 20 mM HEPES pH 7.5, 100 mM sodium chloride and 0.01% MNG/0.001% CHS. The IANBD-amide-labeled minimal cysteine GCGR was concentrated and aliquots were flash frozen for subsequent experiments.

##### Fluorescence experiments

The aliquots of 31C- or 345C-IANBD-amide labeled minimal cysteine GCGR was diluted to 2 µM in 20 mM HEPES pH 7.5, 100 mM sodium chloride, 0.01% MNG/0.001% CHS and 50 µM ZP3780 and incubated at room temperature for 1 hour in order to fully bind agonist. RAMP2 was added to varying fold molar excess with GCGR (0, 0.25, 0.5, 1, 2.5) and incubated at room temperature for 1 hour. Samples were diluted 10-fold with the same buffer for a final GCGR-minimal cysteine-IANBD concentration of 200 nM and final concentrations of RAMP ranging from 0 to 500 nM. Fluorescence data was collected in a 150 µL cuvette with FluorEssence v3.8 software on a Fluorolog instrument (Horiba) in photon counting mode. NBD fluorescence was measured by excitation at 420 nm with excitation and emission bandwidth passes of 5 nm, and the emission spectra was recorded from 505 to 605 nm with 1 nm increments and a 0.5 s integration time. The fluorescence at the peak maximum of 539 nm for all conditions was used for the inset panels Fig. 1E & Fig. 1F as a function of RAMP2 concentration in order to more quantitatively interpret binding of RAMP2 to GCGR.

#### smFRET

##### Preparation of smFRET samples with minimal cysteine GCGR

GCGR-mincys-265C-345C was diluted to 10 µM and incubated with 200 µM RSSR (Enzo Life Sciences) for 2 hours at room temperature. Next the sample was incubated with 14 mM iodoacetamide at RT for 30 minutes followed by 1.5 hours at 4°C. The sample then underwent size exclusion chromatography on a Superdex 200 Increase 10/300 gel filtration column. Monomeric receptors were collected and concentrated to 10 µM and incubated with 100 µM TCEP for 1 hour at room temperature to remove RSSR spin label. The sample was then incubated with 100 µM maleimide LD555 and 100 µM maleimide LD655 for 1 hour at room temperature followed by incubation with 5 mM L-cysteine for 10 minutes followed by removal of free fluorophore, cysteine, and TCEP with size exclusion chromatography, again on a Superdex 200 Increase 10/300 gel filtration column. Aliquots of specifically-labeled, pure protein were flash frozen for later use.

##### Data collection

To inhibit nonspecific protein adsorption, flow cells for single-molecule experiments were prepared as previously described^15^ using mPEG (Laysan Bio) passivated glass coverslips (VWR) and doped with biotin PEG16. Before each experiment, coverslips were incubated with NeutrAvidin (Thermo Fisher), followed by 10 nM biotinylated antibody (mouse anti-FLAG, Jackson ImmunoResearch). Between each conjugation step, the chambers were flushed to remove free reagents. The antibody dilutions and washes were done in T50 buffer (50 mM NaCl, 10 mM Tris, pH 7.5). To achieve sparse immobilization of labeled receptors on the surface, purified labeled receptor was diluted (ranging from 100-fold to 1000-fold dilution) and applied to coverslips. After achieving optimum surface immobilization (~400 molecules in a 2,000 μm^2^ imaging area), unbound receptors were washed out of the flow chamber and the flow cells were then washed extensively (up to 50 times the cell volume).

Receptors were imaged for smFRET in imaging buffer consisting of 3 mM Trolox, 100 mM NaCl, 2 mM CaCl_2_, 20 mM HEPES pH 7.5, 0.01% L-MNG, 0.001% CHS and an oxygen scavenging system (0.8% dextrose, 0.8 mg ml^−1^ glucose oxidase, and 0.02 mg ml^−1^ catalase). All buffers were made in UltraPure distilled water (Invitrogen). Samples were imaged with a 1.65 na X60 objective (Olympus) on a total internal reflection fluorescence microscope with 100 ms time resolution unless stated otherwise. Lasers at 532 nm (Cobolt) and 633 nm (Melles Griot) were used for donor and acceptor excitation, respectively. FRET efficiency was calculated as (I_A_-0.1I_D_)/(I_D_+I_A_), in which I_D_ and I_A_ are the donor and acceptor intensity, respectively, after background subtraction. Imaging was with 100 millisecond acquisition time (10 Hz) with a Photometrics Prime 95B cMOS camera.

##### Data processing

Single-molecule intensity traces showing single-donor and single-acceptor photobleaching with a stable total intensity for longer than 5 seconds were collected (20–30% of total molecules per imaging area). Individual traces were smoothed using a nonlinear filter^16^ with following filter parameters: window = 2, M = 2 and P = 15. Each experiment was performed ≥4 times to ensure reproducibility. smFRET histograms were compiled from ≥100 molecules per condition (100 millisecond time resolution). Error bars in the histograms represent the standard error from ≥4 independent movies. To ensure that traces of different lengths contribute equally, histograms from individual traces were normalized to one before compiling. Gaussian fitting to histograms was done in Origin Pro.

#### Hydrogen-exchange mass spectrometry

##### Hydrogen–deuterium exchange labeling reaction

Prior to HDX, purified RAMP2 in 20 mM HEPES pH 7.5, 100 mM sodium chloride and 0.05% DDM/0.005% CHS, and GCGR in 20 mM HEPES pH 7.5, 100 mM sodium chloride and 0.02% MNG/0.002% CHS, were both separately incubated overnight with PNGaseF to facilitate peptide digestion during LC/MS. Subsequently, 10 µM GCGR was incubated in the presence of saturating RAMP2 (at least 1.5:1 RAMP2:GCGR) at 22 °C for at least one hour. Additionally, 10 µM GCGR was incubated with a volume of 0.05% DDM/0.005% CHS equivalent to the volume of RAMP2 added in preparation of the complex. Similarly, 10 µM RAMP was incubated with a volume of 0.02% LMNG/0.002% CHS equivalent to the volume of GCGR added in preparation of the complex.

To prepare deuterated buffer for HDX, NaCl and HEPES were resuspended in D_2_O (Sigma-Aldrich) to a final concentration of 100 mM NaCl, 20 mM Hepes, pH_read_=7.3. pH_read_ was adjusted with DCl (Sigma-Aldrich) and NaOD (Sigma-Aldrich). To initiate exchange, samples were diluted 1:10 into D_2_O buffer and quenched 1:1 (3 M urea, 20 mM TCEP, pH 2.4), for a total sample volume of 70 µL. At selected time points, 0.5 µL of porcine pepsin (10 mg/ml; Sigma Aldrich) and 0.5 µL of aspergillopepsin (10 mg/ml; Sigma Aldrich) were added to each sample, which was then rapidly vortexed, returned to ice for 3 minutes, and flash frozen in liquid N_2_. Samples were stored at –80 °C prior to LC/MS analysis. Note that proteases were resuspended in 100 mM NaCl, 20 mM HEPES, pH 7.5 to 10 mg/ml and filtered (0.22 µm filter, Corning), aliquoted and stored at –80 °C prior to use.

##### Liquid chromatography/mass spectrometry analysis

Samples were thawed and injected into a cooled valve system (Trajan LEAP) coupled to an LC (Thermo Ultimate 3000) flowing buffer A (0.1% formic acid) at 200 µL/min. Sample time points were injected in non-consecutive order. The valve chamber, trap column, and analytical column were kept at 2 °C.

Peptides were desalted for 4 minutes on a trap column (1 mM ID x 2 cm, IDEx C-128) manually packed with POROS R2 reversed-phase resin (Thermo Scientific). Peptides were then separated on a C8 analytical column (Thermo Scientific BioBasic-8 5 μm particle size 0.5 mM ID x 50 mM) with buffer B (100% acetonitrile, 0.1% formic acid) flowing at a rate of 40 µL/min, increasing from 5% to 50% over the first 26 minutes and from 50% to 90% B over 90 s. The analytical column was washed using two repeating sawtooth gradients and equilibrated at 5% buffer B prior to the next injection. After every fourth injection, a blank injection was performed to monitor for run-to-run carry over. Peptides were eluted directly into a Q Exactive Orbitrap Mass Spectrometer (ThermoFisher) operating in positive ion mode (MS1 settings: resolution 140000, AGC target 3e6, maximum IT 200 ms, scan range 300-1500 m/z). Separate samples of GCGR and of RAMP2 were subject to tandem mass spectrometry analysis (MS1 settings same as above but with resolution 70000, and MS2 settings as follows: resolution 17500, AGC target 2e5, maximum IT 100 ms, loop count 10, isolation window 2.0 m/z, NCE 28, charge state 1 and >8 excluded, dynamic exclusion 15.0 s).

##### Peptide identification and analysis

MS2 data were processed using Byonic (Protein Metrics), which resulted in a list of reference peptides for both GCGR and RAMP2. This reference set included both unmodified sites and de-glycosylated sites within the RAMP and GCGR extracellular domains. Deuterium uptake data were analyzed with HD-Examiner (Version 3.1, Sierra Analytics). HD-Examiner was used with default settings; uptake values were adjusted to account for 1:10 dilution of undeuterated sample into deuterated buffer. For unimodal peaks, changes in deuterium uptake were determined by subtracting the mass centroid of the undeuterated peptide from that of the deuterated peptide. For bimodal peaks, peptides were analyzed using scripts adapted from^17^. Briefly, bimodal peaks were globally fit to sums of Gaussians, such that the widths (and the centers, in the case of bimodals exhibiting EX1 behavior) were constant across time points. Fitting was performed using the *lmfit* package in Python. Deuteration differences for bimodal peptides displayed in the Woods plot (Fig. S4A) were based on the centroids of the far-right (higher mass) peaks, identified using HD-Examiner’s automated bimodal fitting method, when applicable.

